# Inter-membrane association of the Sec and BAM translocons for bacterial outer-membrane biogenesis

**DOI:** 10.1101/589077

**Authors:** Sara Alvira, Daniel W. Watkins, Lucy Troman, William J. Allen, James Lorriman, Gianluca Degliesposti, Eli J. Cohen, Morgan Beeby, Bertram Daum, Vicki A.M. Gold, J. Mark Skehel, Ian Collinson

## Abstract

The outer-membrane of Gram-negative bacteria is critical for surface adhesion, pathogenicity, antibiotic resistance and survival. The major constituent – hydrophobic β-barrel <u>O</u>uter-<u>M</u>embrane <u>P</u>roteins (OMPs) – are secreted across the inner-membrane through the Sec-translocon for delivery to periplasmic chaperones *e.g.* SurA, which prevent aggregation. OMPs are then offloaded to the β-<u>B</u>arrel <u>A</u>ssembly <u>M</u>achinery (BAM) in the outer-membrane for insertion and folding. We show the <u>H</u>olo-<u>T</u>rans<u>L</u>ocon (HTL: an assembly of the protein-channel core-complex SecYEG, the ancillary sub-complex SecDF, and the membrane ‘insertase’ YidC) contacts SurA and BAM through periplasmic domains of SecDF and YidC, ensuring efficient OMP maturation. Our results show the trans-membrane proton-motive-force (PMF) acts at distinct stages of protein secretion: for SecA-driven translocation across the inner-membrane through SecYEG; and to communicate conformational changes *via* SecDF to the BAM machinery. The latter presumably ensures efficient passage of OMPs. These interactions provide insights of inter-membrane organisation, the importance of which is becoming increasingly apparent.

## INTRODUCTION

Outer-membrane biogenesis in Gram-negative bacteria (reviewed in (Konovalova et al., 2017)) requires substantial quantities of protein to be exported; a process which begins by transport across the inner plasma membrane. Precursors of β-barrel <u>O</u>uter-<u>M</u>embrane <u>P</u>roteins (OMPs) with cleavable N-terminal signal-sequences are targeted to the ubiquitous Sec-machinery and driven into the periplasm by the ATPase SecA and the trans-membrane proton-motive-force (PMF) (Brundage et al., 1990; Collinson, 2019; Cranford-Smith and Huber, 2018; Lill et al., 1989). Upon completion, the pre-protein signal-sequence is proteolytically cleaved (Chang et al., 1978; Josefsson and Randall, 1981), releasing the mature unfolded protein into the periplasm. The emergent protein is then picked up by periplasmic chaperones, such as SurA and Skp, which prevent aggregation (McMorran et al., 2013; Sklar et al., 2007), and somehow facilitate delivery to the β-<u>B</u>arrel <u>A</u>ssembly <u>M</u>achinery (BAM) for outer-membrane insertion and folding (Voulhoux et al., 2003; Wu et al., 2005).

In *E. coli,* BAM consists of a membrane protein complex of subunits BamA-E, of known structure (Bakelar et al., 2016; Gu et al., 2016; Iadanza et al., 2016). The core component, BamA, is a 16 stranded β-barrel integral membrane protein, which projects a large periplasmic stretch of 5 <u>PO</u>lypeptide <u>TR</u>anslocation-<u>A</u>ssociated (POTRA) domains into the periplasm. BamB-E are peripheral membrane lipoproteins anchored to the inner leaflet of the OM. Despite structural insights, the mechanism for BAM-facilitated OMP insertion is unknown (Ricci and Silhavy, 2019).

The bacterial periplasm is a challenging environment for unfolded proteins, so complexes spanning both membranes are critical for efficient delivery through many specialised secretion systems (Green and Mecsas, 2016). So, how do enormous quantities of proteins entering the periplasm *via* the general secretory pathway (Sec) efficiently find their way through the cell envelope to the outer-membrane? From where is the energy derived to facilitate these trafficking processes some distance from the energy transducing inner-membrane and in an environment lacking ATP? Could it be achieved by a direct interaction between chaperones, and the translocons of the inner (Sec) and outer (BAM) membranes?

The core-translocon, SecYEG, does not possess periplasmic domains of sufficient size to mediate such an interaction (Van den Berg et al., 2004). However, the <u>H</u>olo-<u>T</u>rans<u>L</u>ocon (HTL) contains the ancillary sub-complex SecDF and the membrane protein ‘insertase’ YidC (Duong and Wickner, 1997; Schulze et al., 2014); both of which contain periplasmic extensions potentially large enough to reach the POTRA domains of BamA.

SecDF is a member of the so called <u>R</u>oot <u>N</u>odulation <u>D</u>ivision (RND) superfamily of proton-motive-force (PMF) driven transporters (reviewed in (Tseng et al., 1999)). It is a highly conserved component of the bacterial Sec translocon, where it has long been known to facilitate protein secretion (Duong and Wickner, 1997; Economou et al., 1995; Pogliano and Beckwith, 1994). While fellow component of the HTL – YidC – is essential for membrane protein insertion, and thus indispensable (Samuelson et al., 2000; Scotti et al., 2000), knock out mutants of *secD* and *secF* are not fatal but severely compromised and cold-sensitive (Gardel et al., 1987), presumably due to deficiencies in envelope biogenesis. The cause of this has been ascribed to a defect in protein transport across the inner membrane.

In keeping with other members of the RND family, like AcrB (Eicher et al., 2014), SecDF confers PMF stimulation of protein secretion (Arkowitz and Wickner, 1994). Different structures of SecDF show the large periplasmic domains in different conformational states (Furukawa et al., 2018; Kumazaki et al., 2014; Mio et al., 2014), also affected by altering a key residue of the proton transport pathway (SecD_D519N_ – *E. coli* numbering) (Furukawa et al., 2017). On this basis, an elaborate mechanism has been proposed whereby PMF driven conformational changes at the outer surface pick up and pull polypeptides as they emerge from the protein channel exit site of SecY. Yet, ATP- and PMF-driven translocation across the inner membrane does not require SecDF or YidC (Gardel et al., 1987; Schulze et al., 2014); SecYEG and SecA will suffice (Brundage et al., 1990). Evidently then, there must be two PMF-dependent components of protein secretion: one early stage dependent on SecYEG/ SecA, and another later event on AcrB-like SecDF activity. This distinction has not been fully considered.

This study explores the role of the ancillary components of the Sec machinery for protein secretion and downstream trafficking through the periplasm for outer-membrane delivery and maturation. In particular, we examine the possibility of a direct interaction between the HTL and BAM machineries to facilitate protein transport through the envelope. The basic properties and structure of the inter-membrane super-complex are investigated, as well as its importance for OMP folding and insertion. The implications of this interaction and its modulation caused by proton transport through SecDF are profound. Thus, we consider their consequences for the mechanism of protein transport through the Sec and BAM machineries, and for outer-membrane biogenesis.

## RESULTS

### Co-fractionation and immunoprecipitation highlight an interaction between the Sec and BAM machineries

Total *E. coli* membranes of cells over-producing either SecYEG or HTL were prepared and fractionated by sucrose gradient centrifugation to separate the inner- and outer-membranes (Fig. 1a). We first sought to determine the precise locations of IM and OM proteins in the fractions; SDS-PAGE analysis of the fractions stained for total protein revealed the presence of SecY in the lighter inner-membrane fractions (Fig. S1a, asterisk – left panel). Heating the fractions (required to unfold outer membrane proteins) prior to SDS-PAGE helped reveal the location of the most highly expressed outer membrane residents (OmpC and OmpF; Fig. S1a, asterisk – right panel). Thus, in these gradients fraction 1-3 mostly contain outer-membranes, and fractions 4-6 are composed mainly of inner-membranes.

**Figure. 1:**
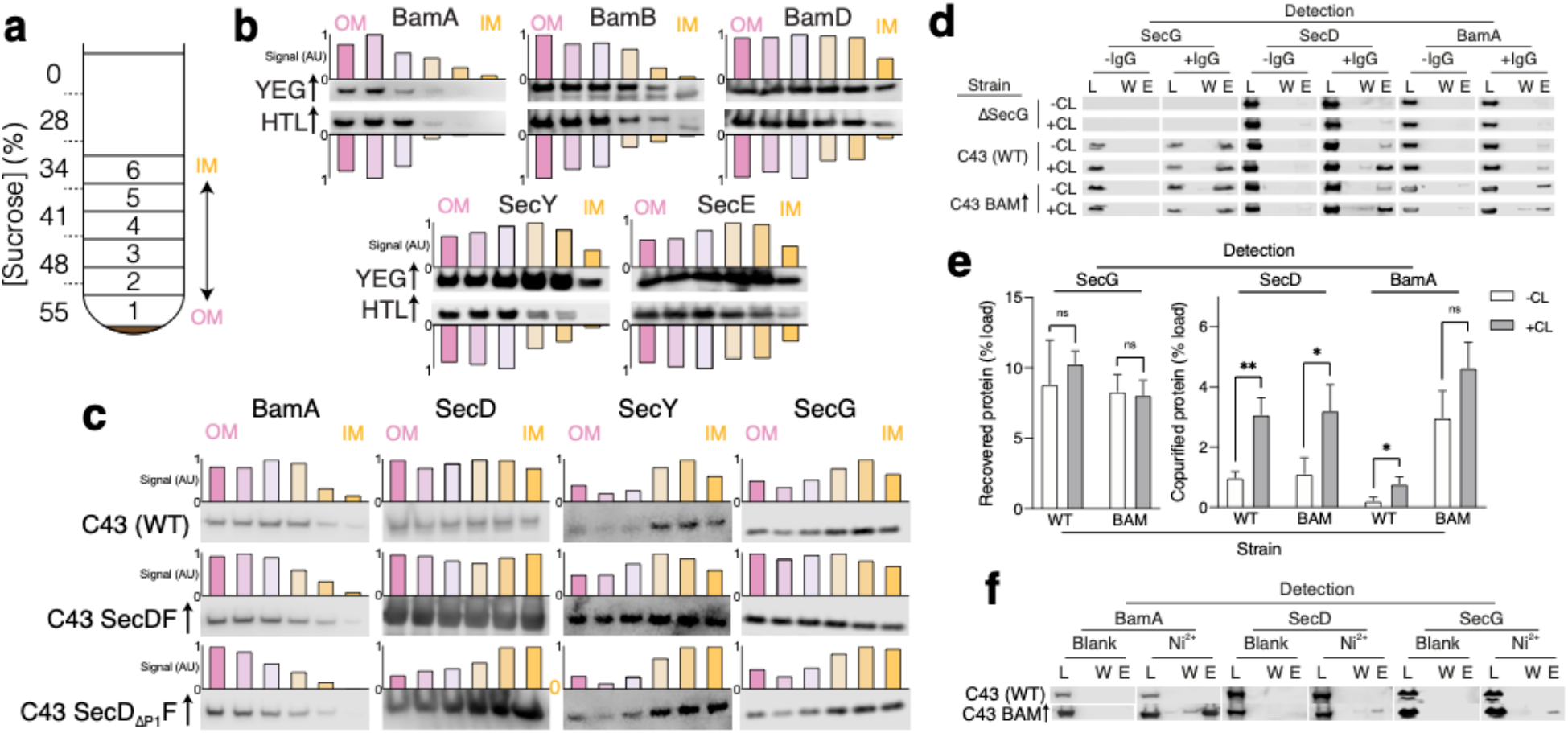
Identification of interactions between HTL and BAM. **a.** Schematic representation of sucrose gradient centrifugation tube for fractionation of *E. coli* total membranes. Numbers 1-6 indicate the fractions taken for SDS PAGE and immunoblotting shown in **b, c** and **d**. **b, c.** Immunoblots of fractions produced as shown in **a** for membranes of *E. coli* C43 overproducing either SecYEG or HTL (**b**) or for *E. coli* C43 with no over-expression, and those over-producing either SecDF or SecD_ΔP1_F (lacking the periplasmic domain 1 (P1) of SecD) (**c**). To help visualise migration shifts, blotting signal was used to quantify relative abundances of proteins of interest, shown above or below blots as normalised bar charts. **d.** Co-immuno-precipitations (co-IP) of SecG, SecD and BamA – pulling with the SecG antibody. Pull-downs were conducted with solubilised crude membrane extracts from *E. coli* C43 (WT), a strain lacking SecG (*ΔsecG*), and C43 over-producing BAM. Experiments were conducted in the presence (+CL) and absence (-CL) of cardiolipin. L = load (1% total material), W = final wash before elution (to demonstrate proper washing of affinity resin, 17% of total material) and E = elution (17% of total material). **e.** Quantification of IPs shown in (**d**). Error bars represent SEM. An unpaired T-test was used to compare samples (p = 0.05, n = 3, * = < 0.05, ** = < 0.01, p values from left to right are 0.4874, 0.8083, 0.0041, 0.0249, 0.0241 and 0.0839). **f.** Affinity pull-down of recombinant BamA-His_6_, SecD and SecG by nickel chelation all in the presence of cardiolipin. L, W and E as described in **(d).**

Immunoblotting confirmed the presence of the BAM components (BamA, BamB and BamD), as expected, in OM fractions (OM; Fig. 1b). Likewise, the over-produced SecY and SecE subunits mark the fractions containing the core-complex (SecYEG) in the IM fractions (IM; Fig. 1b, YEG↑). However, when over-produced as part of HTL there is a marked shift of their migration towards the outer-membrane containing fractions (Fig. 1b, HTL↑). Interestingly, the over-production of SecDF alone results in a similar effect (Fig. 1c); a measurable increase of SecY and SecG in the outer-membrane containing fractions. This effect was lost in comparable experiments where the periplasmic domain of SecD (P1) had been removed (Fig. 1c). Our interpretation of these experiments is that the interaction between the Sec and Bam complexes, requiring at least the periplasmic domains of SecD (and most likely SecF and YidC), causes an association of inner and outer membrane vesicles reflected in the shift we observe.

To further examine this interaction, we extracted native membranes with a mild detergent for <u>I</u>mmuno-<u>P</u>recipitation (IP) using a monoclonal antibody raised against SecG. The pull-downs were then probed for native interacting partners by western blotting (Fig. 1d,e, Fig. S1b). As expected, SecG (positive control) and SecD of HTL co-immuno-precipitated. Crucially, BamA could also be detected. The specificity of the association was demonstrated by controls omitting the SecG antibody or the SecG protein (produced from membranes extracts of a *Δ*secG strain (Nishiyama et al., 1994)), wherein non-specific binding was either undetectable, or considerably lower than the specific co-immuno-precipitant (Fig. 1d). When BAM was over-produced, the yield of BamA recovered in the IPs increased accordingly (Fig. 1d, e, Fig. S1b).

In a similar experiment, a hexa-histidine tagged BamA was used to isolate BAM from cells over-producing the complex. Western blots showed that BamA co-purified, as expected, with additional components of the BAM complex (BamB and BamD), and crucially also with SecD and SecG of the HTL (Fig. 1f, Fig. S1c). Again, controls (omitting Ni^2+^, or recombinant his_6_-BamA) were reassuringly negative.

### Interaction between HTL and BAM is cardiolipin dependent

The phospholipid <u>C</u>ardio<u>L</u>ipin (CL) is known to be intimately associated with energy transducing systems, including the Sec-machinery, for both complex stabilisation and efficient transport (Corey et al., 2018; Gold et al., 2010; Schulze et al., 2014). For this reason, the IP experiments above were augmented with CL. On omission of CL the interactions of SecG with SecD and BamA were reduced ~ 3- and 5-fold, respectively (Fig. 1d, e; Fig. S1b). This lipid-mediated enhancement of the SecG-SecD interaction is consistent with our previous finding that CL stabilises HTL (Schulze et al., 2014), and shows it also holds true for the HTL-BAM interaction. *Apropos*, CL has been shown to be associated with the BAM complex (Chorev et al., 2018).

### HTL and BAM interact to form an assembly large enough to bridge the inner and outer membranes

To confirm the interaction between the Sec and BAM machineries, the purified complexes were subjected to glycerol gradient centrifugation. When mixed together, HTL and BAM co-migrated towards higher glycerol concentrations, beyond those attained by the individual complexes (Fig. 2a); consistent with the formation of a larger complex due to an interaction between the two. The interaction is clear, but not very strong, likely due to the required transient nature of the association between the two translocons *in vivo*, and also because of the complete breakdown of the inner and outer membranes by detergent – required for this experiment. When the experiment was repeated with the individual constituents of HTL: SecDF and YidC, but not SecYEG, were also shown to interact with BAM (Fig. S2a-c).

**Figure. 2:**
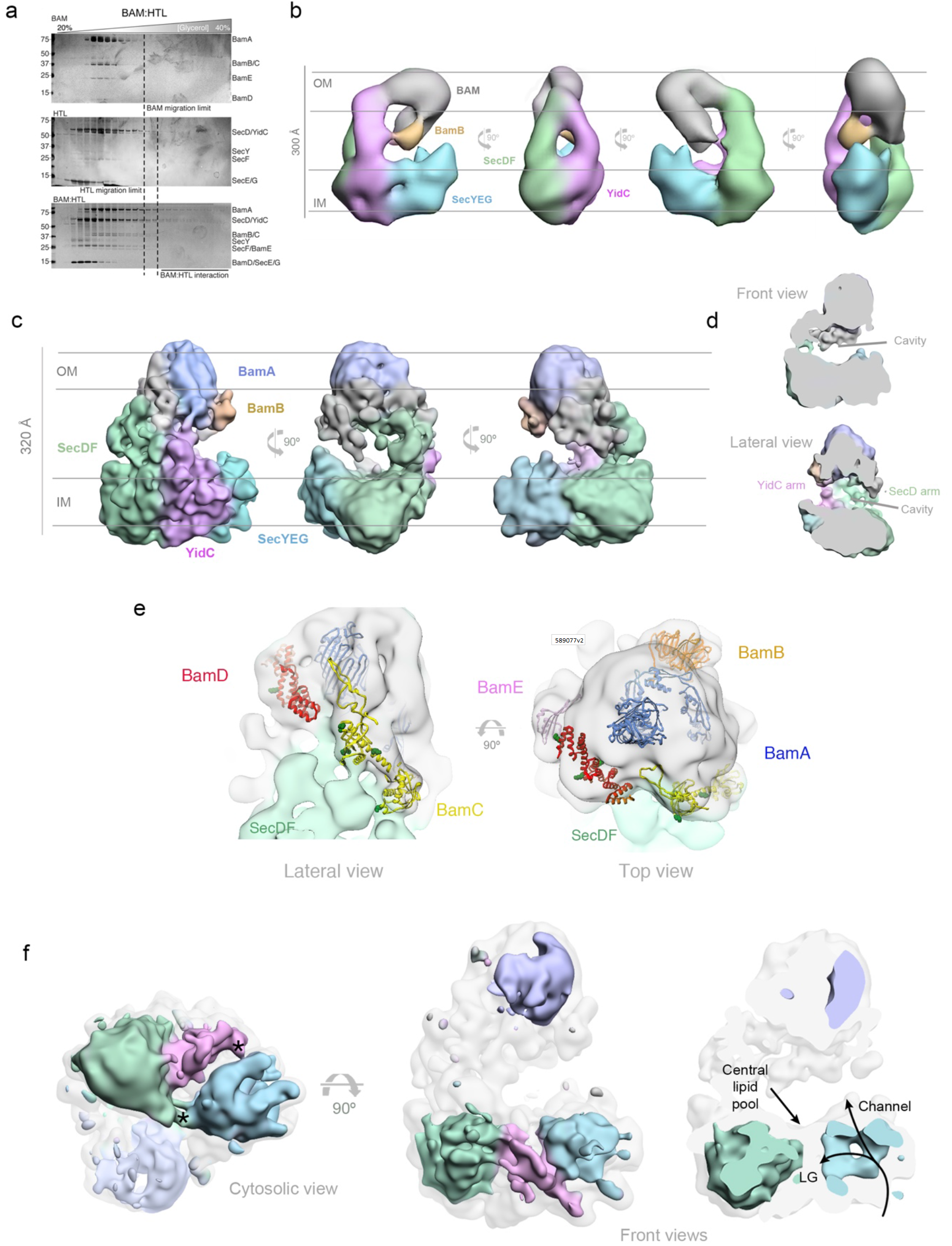
3D characterization of HTL-BAM by NS-EM and cryo-EM in detergent solution, and XL-MS analysis. **a.** Silver-stained SDS-PAGE gels of fractions from glycerol centrifugation gradients are shown, with increasingly large complexes appearing in fractions of higher percentage glycerol, (from left to right). The glycerol centrifugation gradients of BAM alone (top) HTL alone (middle) and HTL mixed with BAM (bottom) are shown. Dashed lines represent the fraction of furthest migration of the individual components, as determined in the top two gels. **b.** Negative stain analysis of the HTL-BAM complex (37.2 Å resolution) in four representative orthogonal views, with the orientation with respect to the inner and outer membranes inferred. BAM (grey), BamB (orange), SecYEG (cyan), YidC (pink) and SecDF (green) are shown. **c.** Three orthogonal views of the cryo-EM HTL-BAM complex 3D reconstruction (27 Å resolution). Colours are as in (**a)**, but with BamA in blue. **d.** Frontal and lateral sliced views of the cryo HTL-BAM complex showing the large cavity between the IM and OM complexes. **e.** Close-up of the outer-membrane region of the HTL-BAM complex. The cryo-EM structure (transparent surface) with BamABCDE atomic structures docked (pdb_5d0q). BamB (orange) position was determined directly by NS-EM (Fig. S4e). BamA (blue), BamC (yellow), BamD (red) and BamE (pink) are docked according to the HTL-BAM cryo-EM density and XLMS data (Fig. S6c). Green sphere atoms in BamC and BamD show interacting points with SecD identified by mass apectrometry. **f.** Lower threshold map of HTL-BAM overlaid with the correct level (transparent grey), with the main components coloured as in (**a**). The lateral gate (LG) into the membrane, protein-channel through SecY and the central lipid pool are highlighted. The asterisks denote interaction sites between SecYEG and SecDF/ YidC.

Visualisation of the heavy fractions containing interacting HTL and BAM by Negative Stain (NS)-Electron Microscopy (EM) revealed a heterogeneous mixture of small and very large complexes (Fig. S3a, large complexes marked with white arrows). As noted above, this mixed population is probably due to the expected transient nature of the interaction between the two complexes, and/ or due to super-complex instability caused by loss of the bilayer and specifically bound phospholipids, *e.g.* CL, during purification (see above and below). Even though we augment the material with CL it is unlikely the full complement of native lipids found in the membrane bound state are restored.

To overcome this heterogeneity we stabilised the complex by cross-linking, using GraFix (Kastner et al., 2008) (Fig. S4a, middle). Note that this interaction was also demonstrated by size exclusion chromatography, performed for sample preparation for mass spectrometry and cryo-EM (see below). Visualisation by NS-EM revealed a marked reduction in the number of dissociated complexes (Fig. S3b). As expected, omitting CL from the preparation resulted in dissociation of the majority of the large complexes, even with GraFix (Fig. S3c), supporting the findings above of the CL dependence of the interaction (Fig. 1e).

Mass spectrometry confirmed the presence of BAM and HTL constituents in a high molecular weight GraFix fraction (Fig. S4a, right; Table S1), which was selected for analysis by NS-EM. The subsequent single particle analysis of this material (Table S2) revealed a remarkable structure large enough (~ 300 x 250 x 150 Å) to contain both Sec and BAM machineries (Botte et al., 2016; Iadanza et al., 2016), and with a height sufficient to straddle the space between the two membranes (Fig. 2b, Fig. S4b), especially when considering the plasticity of the periplasm (Zuber et al., 2008). Moreover, the periplasmic domains of the HTL and BAM complexes are potentially large enough reach out across the space between the inner and outer-membranes to contact one another. Indeed, regions of SecDF and the POTRA domains of BamA have been show to extend ~ 60 Å (Furukawa et al., 2017) and ~ 110 Å (Ma et al., 2019) respectively, sufficient to bridge this gap.

To assign the locations and orientations of the individual constituents of HTL and BAM, we compared the 3D reconstructions of different sub-complexes: BAM bound to SecYEG-DF (without YidC) (Fig. S4c) or SecDF alone (Fig. S4d). The difference analysis revealed the locations of YidC (Fig. 2b, pink; Fig. S4c, pink arrow), SecDF (Fig. 2b, green; Fig. S4d, green arrow) and SecYEG (Fig. 2b, blue; Fig. S4d, blue arrow) at the bottom of the assembly (assigned as the inner-membrane region). The orientation of BAM relative to SecDF is different in SecDF-BAM compared to HTL-BAM (Fig. S4d, red arrows), possibly due to its known ability to move (see below), and/ or the absence of stabilising interactions with the rest of HTL components.

Removing BamB from the complex results in the loss of significant mass in the area designated as the OM region with respect to the holo-structure (Fig. 2b, orange; Fig. S4e, orange arrow). This confirmed the orientation of the inner and outer-membranes, and the assignment of the BAM complex as shown in Fig. 2b. Interestingly, the complex lacking BamB shows a diminishment of the density assigned as YidC (Fig. S4e, pink arrow), suggestive of a mutual interaction between the two.

### Periplasmic domains of the Sec and BAM translocons associate to form a large cavity between the bacterial inner and outer membranes

Despite heterogeneity in the sample we were able to isolate a cross-linked HTL-BAM complex by size exclusion chromatography and produce a low-resolution cryo-EM structure (Fig. 2c, Fig. S5 and Table S2) with an overall resolution of 18.23 Å.Taken together with the difference map generated by NS-EM (Fig. 2b; Fig. S4) the structure reveals the basic architecture of the assembly and the arrangement of constituent subunits.

The extreme complexity of the image processing resulted in an insuffient number of particles to attain high resolution. Many factors contribute to this problem: the dynamism of the complex, due to the limited contact surface between the HTL and BAM; its inherent mobility necessary for function; the presence of detergent surrounding the trans-membrane regions of the HTL and BAM components, accounting for most of the surface of the assembly; and finally, the absence of inner- and outer-membranes scaffolds. The loss of the fixed double membrane architecture was particularly problematic; during image processing we found different sub-populations where the BAM pivots away from its raised position towards where the inner membrane would otherwise have been. Obviously, this would not happen if restrained by the outer-membrane.

In spite of all this, the attainable structure proved to be very illuminating. Due to the limited resolution, we deployed cross-linking mass spectroscopy (XL-MS) to verify the contacts between HTL and BAM responsible for inter-membrane contact points. The HTL and BAM complexes were mixed together in equimolar quantities and cross-linked with the lysine-specific reagent DSBU. The reaction mixture was then fractionated by gel filtration and analysed by SDS-PAGE. A single band corresponding to the cross-linked HTL-BAM complex was detected (Fig. S6a); note that the isolation of the intact HTL-BAM complex by size exclusion chromatography provides further evidence of a genuine interaction. The fractions containing the cross-linked complex were combined and digested before LC-MS/ MS analysis. The analysis of mass spectrometry data enabled the detection and mapping of the inter- and intra-molecular protein cross-links within the assembly. The results show an intricate network of interactions most of which are consistent with the cryo-EM structure, particularly at one side of the assembly between SecD and BamBCD, and on the other side between YidC and BamABCD (Fig. S6b,c). All the constituent proteins of HTL were cross-linked to BAM subunits with exception of SecG and YajC. Thus, the resulting co-immunoprecipitation and affinity pull down of SecG with the Bam components (described above; Fig. 1d-f) must have been the result of an indirect interaction, presumably *via* SecDF-YidC. This is consistent with the lack of an interaction of SecYEG alone and the BAM complex (Fig. S2b), and the assignment of the electron microscopy structures (Fig. 2b,c) – showing no connection between SecYEG and BAM. In this respect, it is interesting to note in the structure that the periplasmic domains of SecD, YidC and to a lesser degree SecF, extend to establish multiple interactions with the BAM lipoproteins suggesting a pivotal role for these subunits in the formation of the HTL:BAM complex (Fig. 2c,e). This bridge between the two complexes also helps to define a very large cavity between the IM and OM (Fig. 2d).

The BAM complex is recognisable in the cryo-EM structure at the outer membrane with the expected extensive periplasmic protrusions (Bakelar et al., 2016; Gu et al., 2016). Some components of the BAM complex, such as BamB, can be unambiguoulsy docked at the cryo-EM structure, due to the direct determination by NS and its easily recognisable ß-propeller shape (Fig. 2e). We suggest also the locations of BamA, BamC and BamD according to the cryo-EM density and the constraints of the XL-MS data (Fig. 2e; Fig. S6c).

The inner-membrane region the HTL is much more open than the previous structure, when visualised alone (Botte et al., 2016). In the new open structure, the locations of the core-complex SecYEG, SecDF and YidC can be easily distinguished, connected by two bridges from the former (Fig. 2f, left, asterisks). These bridging sites could be the binding sites of CL and the central lipid pool identified previously, required for structural stability and translocon activity (Corey et al., 2018; Martin et al., 2019; Schulze et al., 2014). Within SecYEG the protein channel can be visualised though the centre, along with the lateral gate (required for signal sequence binding and inner membrane protein insertion) facing towards SecDF, YidC and the putative central lipid pool (Martin et al., 2019; Fig. 2f, right).

### Cardiolipin, required for super-complex formation, stabilises an ‘open’ form of the HTL

The HTL bound to BAM in our EM structure (Fig. 3a, structure ii) seems to be more open when compared to the previously published low-resolution cryo-EM structure (Botte et al., 2016) (emd3056; Fig. 3a, structure i), and also displays a more prominent periplasmic region. Preparations of HTL alone, made in this study, contain both a ‘compact’ state (Fig. 3a, structure iii) similar to that of the previously published structure (Fig. 3a, structure i), as well as a proportion of an ‘open’ state, with proud periplasmic domains, not previously described (Fig. 3a, structure iv), and apparently more similar to that seen in the HTL-BAM structure (Fig. 3a, structure ii).

**Figure 3:**
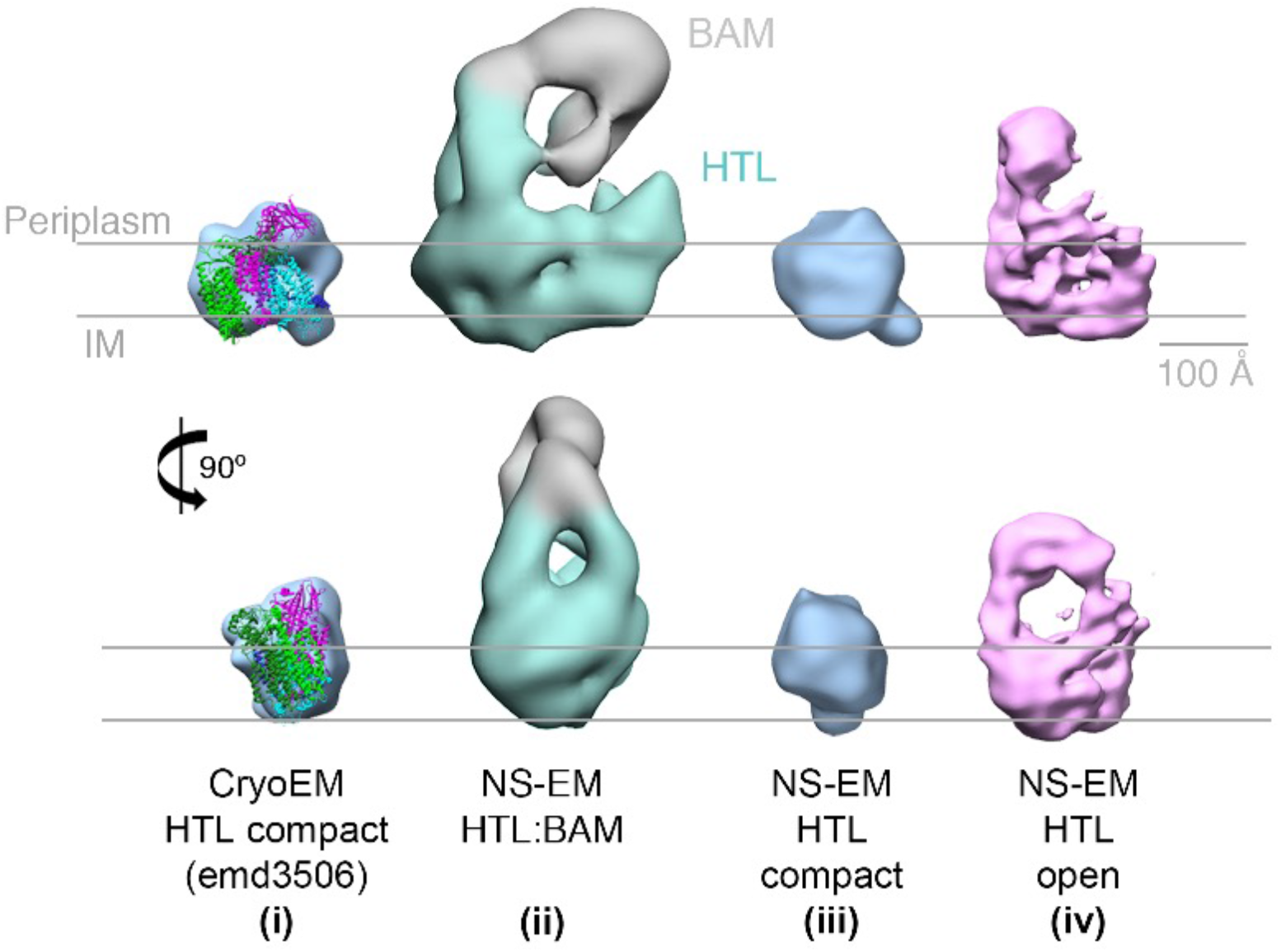
EM structures of HTL in ‘compact’ and ‘open’ states. Structure and docking of a previously published cryo-EM structure of HTL in the compact state **(i)**(Botte et al., 2016), the HTL-BAM complex **(ii)**, HTL in the ‘compact’ state **(iii)** and HTL in the ‘open’ state **(iv)**; structures **(ii - iv)** are from this study.

The HTL sample used here is extremely pure, of known subunit composition and not prone to oligomerisation (Schulze et al., 2014). So, we can rule out that this larger form, assigned as an ‘open’ state, of HTL is not due to the presence of contaminants, unknown additional partner proteins, or dimerisation. Instead, the reason for the presence of these different populations – ‘compact’ and ‘open’ states – is likely due to varying interactions with lipids, including CL. Lipid content within the HTL is critical for proper structure and function, and CL is particularly important for protein translocation through the Sec machinery (Corey et al., 2018; Gold et al., 2010; Hendrick and Wickner, 1991; Schulze et al., 2014). Depletion of these core lipids, for instance by detergent extraction, might be expected to cause a collapse of the complex. Augmenting the HTL with CL during purification increased the proportion of the ‘open’ state (from 8% to 17%), which could be enriched by glycerol gradient fractionation (to 32%), and further stabilised by cross-linking (to 40%) (Fig. S7).

Evidently then, it seems likely that the open conformation (Fig. 3a, structure iv) is the state capable of interacting with the BAM complex (Fig. 3a, structure ii). The ‘open’ structure, and the ‘compact’ structure seen before (Botte et al., 2016), may reflect different functional states of the translocon. Presumably, the HTL would be closed when idle in the membrane, and would open to various degrees depending on the associated cytosolic partners (*e.g.* ribosomes or SecA), periplasmic factors (chaperones, BAM, *etc*.) and various substrates. Thus, it is not suprising that when free of the constraints of the membrane, and in the harsh environment of a detergent micelle, that these various states can be adopted.

### Increasing the distance between inner and outer membranes weakens the HTL-BAM interaction

The dimensions of the HTL-BAM structure roughly corresponds to the distance between the inner and outer membranes. Increasing the thickness of the periplasm might therefore be expected to reduce formation of HTL-BAM complexes, as previously observed for other trans-periplasmic complexes (Asmar et al., 2017; Cohen et al., 2017). To test this prediction, we increased the thickness of the periplasm by manipulating the width-determining lipoprotein Lpp, which separates the outer membrane from the peptidoglycan layer. Increasing the length of *lpp* thus increases the width of the periplasm, from ~ 250 Å for wild-type *lpp* to ~ 290 Å when an additional 21 residues are added to the resultant protein (Lpp_+21_) (Asmar et al., 2017) (Fig. 4a).

**Figure 4:**
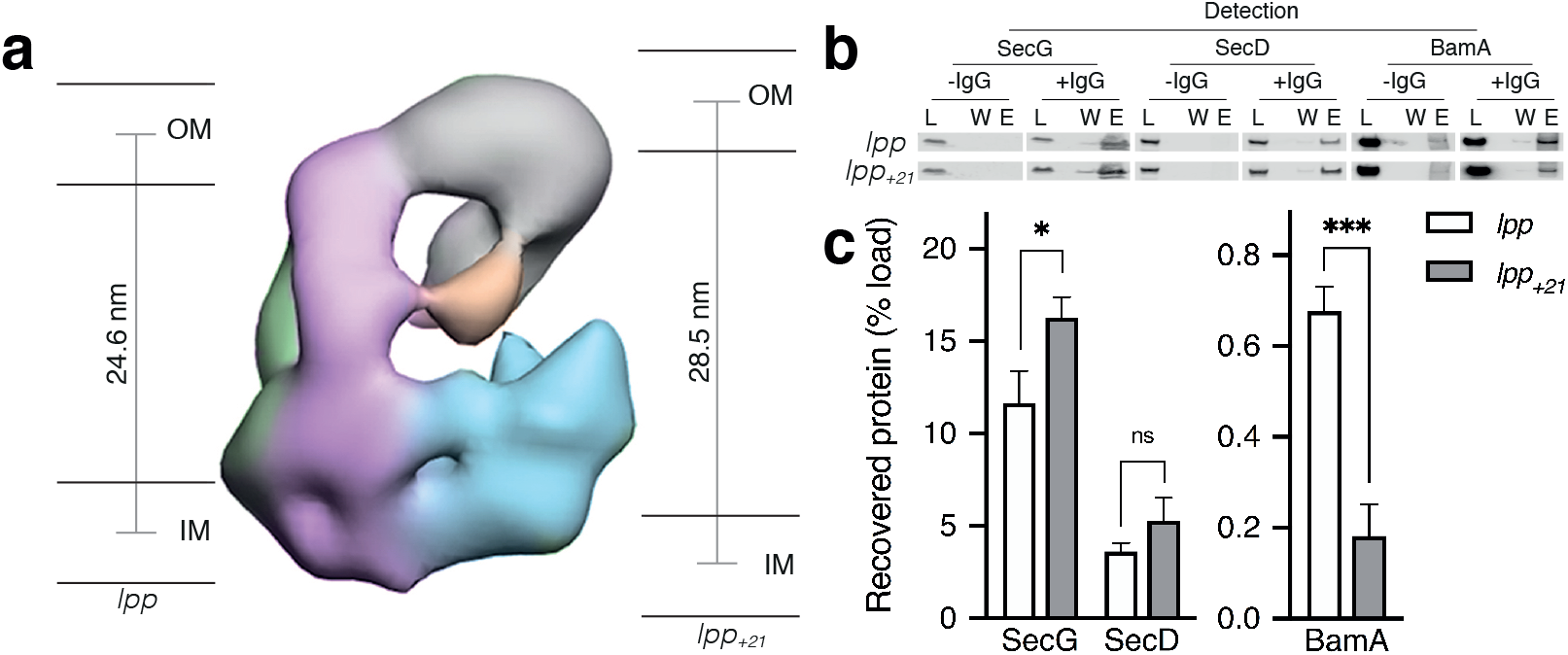
Effect of increasing the periplasmic distance on the HTL-BAM interaction. **a.** Negative-stain EM model of HTL-BAM (from Fig. 2a), annotated with membranes at the experimentally determined distances between the inner and outer membranes of *E. coli* strains containing wild type *lpp* and mutant *lpp_+21_* (Asmar et al., 2017). **b.** Co-immunoprecipitation of SecG, SecD and BamA when pulling from an anti-SecG monoclonal antibody. Co-IPs were conducted in the presence of cardiolipin as in Fig. 1d, but with solubilised membranes of strains described in **(a).** **c.** Quantification of IPs from **(b)**. Error bars represent SEM. An unpaired T-test was used to compare samples (p = 0.05, n = 3, * = < 0.05, *** = < 0.001, p values from left to right are 0.0170, 0.0990 and 0.0006).

The experiments described above (Fig. 1d,e) were repeated: extracting total membranes in the presence of CL for IP by antibodies raised against SecG. Blotting for SecD and BamA then provided a measure respectively for interactions between HTL and BAM (Fig. 4b,c, Fig. S8) in the *lpp_+21_* background. Consistent with our model, the integrity of the HTL in the inner membrane was unaffected, but the recovery of HTL-BAM was reduced more than 3-fold.

### PMF stimulation of protein translocation through the inner membrane by SecA and SecYEG is not conferred by proton passage through SecD

It has been known for many years that SecDF plays a critical role in protein secretion. The results above show that the periplasmic domains of HTL, and in particular those of SecDF, mediate the recruitment of the BAM complex, likely to facilitate the onward journey of proteins to the outer membrane. Therefore, we decided to re-evaluate the precise role and activity of this ancillary sub-complex. Experiments were established to investigate: (1) the role of SecDF in SecA dependent protein transport through the inner membrane *via* SecYEG, and (2) the consequences of its interaction with the BAM machinery for outer membrane protein maturation. In particular, we set out to explore the possibility of an active role in these events for the proton translocating activity of the SecDF sub-complex.

*secDF* null mutants exhibit a severe export defect, and are only just viable (Pogliano and Beckwith, 1994). To explore this phenotype further we utilised *E. coli* strain JP325, wherein the production of SecDF is under the control of an *ara* promoter: the presence of arabinose or glucose, respectively results in production or depletion of SecDF-YajC (Economou et al., 1995) (Fig. 5a, Fig. S9a). When plated, depletion of SecDF-YajC results in a strong growth defect (Fig. 5b, panels 1 and 2), which can be rescued by recombinant expression of wild type *secDF* (Nouwen and Driessen, 2002) (Fig. 5b, panels 3 and 4). In contrast, expression of *SecD_D519NF_*, which results in the production of a complex defective in proton transport (Furukawa et al., 2017), did not complement the defect (Fig. 5b, panels 5 and 6).

**Figure 5:**
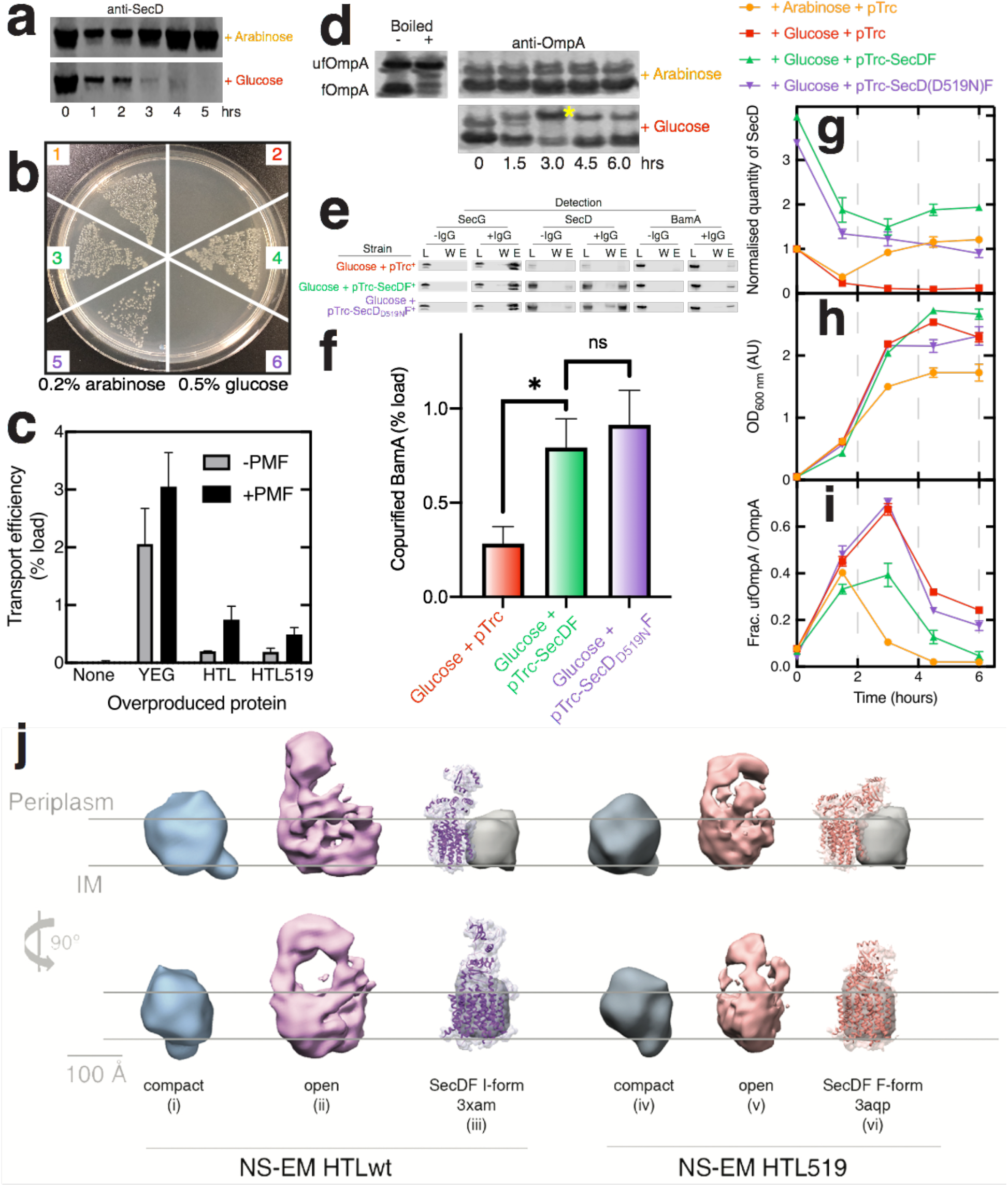
Effects of SecD depletion upon cell growth, OmpA secretion and maturation. **a.** Western blot illustrating depletion of SecD in *E. coli* JP325 whole cells when grown in the presence of arabinose or glucose. t=0 represents the time at which an overnight culture (grown in arabinose) was used to inoculate a fresh culture containing either arabinose or glucose. **b.** Growth of *E. coli* JP325 transformed with empty vector pTrc99a (1 + 2), pTrc99a-*secDF* (3 + 4) and pTrc99A-*SecD_D519NF_* (5 + 6). Samples in panels 1, 3 and 5 were grown on LB-arabinose (permissive), whereas 2, 4 and 6 were grown on LB-glucose (non-permissive). **c.** Classical SecA-driven import assay with *E. coli* inner membrane vesicles (IMVs) and OmpA. IMVs contained over-produced protein as stated on the x-axis. Error bars represent SEM (n=3). **d.** Periplasmic fractions of *E. coli* JP325 immunoblotted for OmpA. Two species of OmpA are visible: folded OmpA (bottom band; fOmpA) and unfolded OmpA (top band; ufOmpA). Also shown are control lanes containing *E. coli* whole cells with over-produced, mainly ‘folded’ OmpA (fOmpA, bottom band) and the same sample, but boiled, to produce ‘unfolded’ OmpA (ufOmpA, top band). **e.** Representative western blots of co-immuno-precipitations conducted as in Fig. 1d in the presence of CL, but with solubilised membranes prepared from *E. coli JP325* grown in the presence of glucose and cloned with variants of pTrc99a, as stated in the figure. **f.** Quantification of BamA pull down from co-IPs shown in **(e)**. Error bars represent SEM. An unpaired T-test was used to compare samples (p = 0.05, n = 3, * = < 0.05, p values from left to right are 0.0449 and 0.6412). For **(g-i)**, samples were prepared from various cell cultures; see key (above) for strains used. Error bars represent SEM (n=3 for experimental samples grown in glucose). **g.** Quantification of SecD from western blots such as those shown in **(a)**. Values are normalised to JP325-pTrc99a at t = 0. **h.** Culture growth curves. **i.** Analysis of western blots such as those from **(d)** showing the quantity of ufOmpA as a fraction of the total OmpA in the periplasmic fraction. **j.** Comparison of NS-EM structures of ‘compact’ (structures i and iv) and ‘open’ (structures ii and v) conformations of HTL *versus* the counterpart containing SecD_D519N_F (HTL519); both in the presence of CL. Atomic structures of SecDF overlaid with filtered maps at 5 Å are shown alongside for the *I*-form (structure iii, 3XAM) and *F*-form (structure vi, 3AQP), with the amino acid substitution equivalent to the *E.coli* SecD_D519N_, 3AQP). The grey arbitrary mass indicates the approximate position and mass of SecYEG.

In light of these findings, we set up a classical *in vitro* transport assay investigating SecA-driven proOmpA transport into inner membrane vesicles (IMVs) containing either over-produced native HTL, or the defective version of HTL (containing SecD_D519N_F). Both sets of vesicles contained similar concentrations of SecY (Fig. S9b), yet despite the blocked proton pathway through SecDF, there was no difference in the efficiencies of transport (Fig. 5c). The lower quantities of transported pre-protein compared to the core-complex (SecYEG) most likely reflects the reduced quantities of SecYEG in the HTL experiments, measured by blotting for SecY (Fig. S9b).

Most importantly, the results demonstrate that HTL does not require a functional proton wire through SecDF for SecA mediated ATP and PMF driven protein translocation (Fig. 5c). In this respect, SecYEG and SecA are sufficient (Brundage et al., 1990). Therefore, the proton translocating activity of SecD, crucial for cell survival, must be required for something else other than secretion through the SecYEG core complex.

### Interaction between the Sec and BAM complexes is required for efficient OmpA folding

The most obvious function of an interaction between the Sec and BAM machineries would be to facilitate efficient delivery and insertion of OMPs to the OM. We therefore reasoned that disrupting this interaction might compromise OMP delivery to BAM, leading to the accumulation of unfolded OMPs in the periplasm – particularly where high levels of OM biogenesis are required, such as in rapidly dividing cells. Elevated levels of unfolded OmpA (ufOmpA) in the periplasm are a classical signature of OMP maturation deficiencies (Bulieris et al., 2003; Sklar et al., 2007), and can be easily monitored by SDS-PAGE and western blotting: folded OmpA (fOmpA) does not denature fully in SDS unless boiled, it therefore runs at a lower apparent molecular mass on a SDS-PAGE compared to ufOmpA (Fig. 5d, left) (Bulieris et al., 2003; Sklar et al., 2007).

Based on the above results, SecDF looks like the most important mediator of the Sec-BAM interaction. We therefore used the SecDF depletion strain (JP325) as a basis for functional assays. To overcome the growth defect (Fig. 5b, panels 1 and 2) and produce sufficient cells to analyse, overnight cultures of the strains used above were grown in permissive media (arabinose), and then transferred to new media containing glucose. At OD_600nm_ = 1.0, the cultures were harvested and membranes were prepared and solubilised for analysis by co-IP as described above. As expected, depletion of SecD (Fig. S9c-d) reduced pull down of BamA (approximately 3-fold) compared to the complemented cells (Fig. 5e-f; Fig. S9c-d).

Overnight cultures were next prepared as described above, and then transferred to new media containing either arabinose or glucose, marked as t = 0 in Fig. 5a, d, g-i. Samples were then taken at regular intervals and the ratio of unfolded to folded OmpA determined (Fig. 5d, i), along with cell density (Fig. 5h) and SecD levels (Fig. 5g). Under SecDF depletion conditions (red squares), high levels of unfolded OmpA accumulate in the periplasm, particularly during the exponential phase where the demand for OM biogenesis is very high (Fig. 5d, asterisk; Fig. 5h,i; Fig. S9e). Meanwhile, under permissive conditions (Fig. 5g-i, arabinose, yellow circles), an increase in ufOmpA is observable after 1.5 h, but it recovers fully by 3 h. Notably, this change is accompanied by a transient decrease in SecDF levels.

Clearly, the expression of *secDF* and levels of ufOmpA in the cell envelope are anti-correlated, exacerbated during fast cell growth. These effects were not an indirect consequence of BamA loss, which was unperturbed (Fig. S9f). Taken together, the data show that depletion of SecDF reduces the interaction between HTL and BAM, and thereby hampering transport of β-barrel proteins to the outer-membrane – resulting in a build-up of ufOmpA in the periplasm. This would compromise the integrity of the outer-membrane and explain the cold-sensitivity of *secDF* mutants (Gardel et al., 1987).

### Proton transport through SecD is required for efficient outer membrane protein maturation

Proton translocation through SecD is crucial for cell growth (Fig. 5b, panels 5 and 6), but evidently not for PMF-stimulated transport through the inner membrane *via* SecYEG (Fig. 5c). To determine if this activity is required for downstream events, we once again deployed the SecDF depletion strain, complemented with wild-type or mutant *secDF* (as above, Fig. 5b, panels 3 - 6). Comparable quantities of the respective SecD variants could be produced (Fig. 5g, green and purple; Fig. S9a). The defective HTL(SecD_D519N_F) protein retained its ability to interact with the BAM complex (Fig. 5f; Fig. S9c,d). The analysis showed the wild type, but not the mutant, reduced unfolded OmpA in the periplasm to levels much closer to that of the strain grown in permissive conditions (Fig. 5i; green and purple, respectively; Fig. S9e). Therefore, proton transport through SecDF is apparently required for efficient outer membrane protein folding.

### HTL(SecD_D519N_F) adopts a different conformation to the native version

The PMF-dependent mobility of the periplasmic domain of SecD (Furukawa et al., 2017) may be critical for its activity as part of the BAM-HTL complex. To test this, the variant of HTL containing SecD_D519N_ was produced for comparison with the native form. Electron microscopy was used to assess the extent of the ‘compact’ and ‘open’ forms of the HTL complex (Fig. 5j, respectively structures i and ii). The 2D classification of HTL-SecD_D519N_ shows the open state is populated to a similar extent compared to the unmodified HTL (Fig. S10).

The 3D analysis shows the compact states in both cases, similar to those seen before (Botte et al., 2016) (Fig. 3, structure i; Fig. 5j, structures i and iv). However, the ‘open’ states are significantly different: blocking the proton pathway in SecD results in a shorter extension of the periplasmic domains of the HTL, compared to the unmodified form (Fig. 5j, structures ii *versus* v). This is consistent with the conformational change observed at atomic resolution in SecDF alone (Fig. 5j, structures iii *versus* vi) (Furukawa et al., 2017). Even at the current low-resolution description of the HTL-BAM complex (Fig. 2), it is clear that these observed PMF-dependent conformational changes of SecD would be communicated to the outer membrane.

### HTL interacts with the periplasmic chaperone SurA in complex with OmpA

The results above lead us to address outstanding questions regarding chaperone interaction with the Sec complex. When proteins emerge from the channel into the periplasm they are presumably quickly engaged by chaperones, prime examples being Skp and SurA (McMorran et al., 2013; Sklar et al., 2007). So far, no evidence has been presented of a direct interaction of these chaperones with the Sec-machinery. As noted above, the core SecYEG complex does not contain periplasmic domains large enough to do so. We explored the possibility of such an interaction with the HTL, using the same approach described above for HTL variants and BAM (Fig. 2a, Fig. S4a).

Addition of SurA to the HTL (without cross-linking) followed by glycerol gradient centrifugation yielded heavy fractions containing HTL and SurA (Fig. S11a), indicative of them being in complex. As in the case for the HTL-BAM binding experiment, only low yields of the HTL-SurA complex were generated. Again, this is understandable because of the likely transient nature of the translocon-chaperone interaction. To overcome this the experiment was repeated in the presence of cross-linker (Grafix) and fractions were prepared for NS-EM (Fig. S11b).

The subsequent analysis revealed a structure with dimensions of roughly 300 x 250 x 150 Å, large enough to contain both HTL and SurA (Fig. 6a). To overcome the uncertainty, due to the low yield of HTL-SurA recovery, we decided to verify the presence and location of SurA on the isolated complex. To do so, the complex was immuno-decorated with a polyclonal antibody raised against SurA (Fig. 6b. Fig. S11c-d) or, as a control, a monoclonal antibody specific to a cytosolic loop of SecY (Corey et al., 2016) (Fig. 6c, Fig. S11e-f). The results mark the location of the cytosolic face of SecY (Fig. 6c), and SurA as a prominent extention on the other periplasmic side of the inner-membrane closest to SecDF-YidC.

**Figure. 6:**
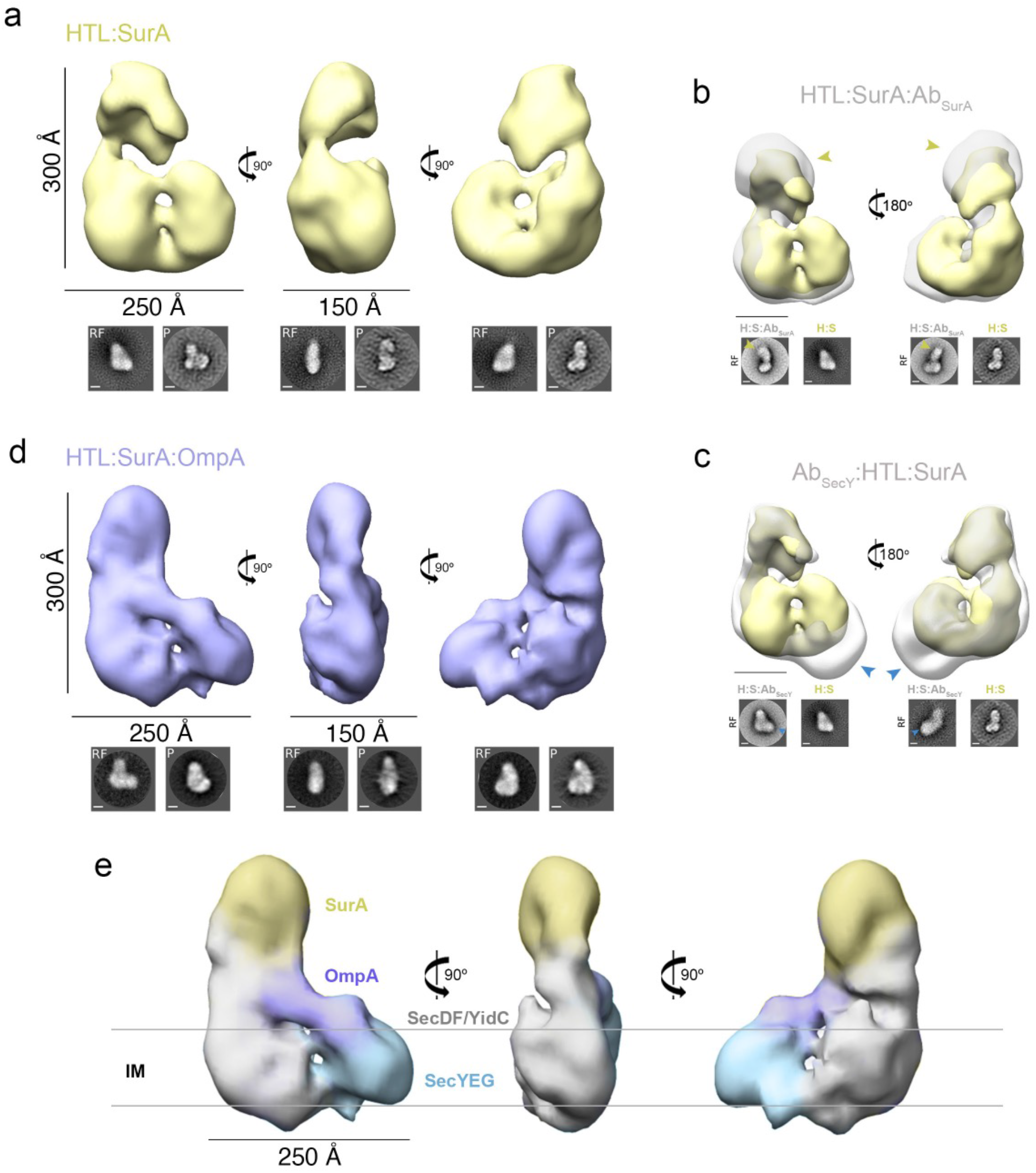
Negative-stain analysis of HTL:SurA and HTL:SurA-OmpA complexes. For **a**, **b, c** and **d** top panel shows views of 3D reconstructions, and bottom shows reference-free (RF) class averages and projections (P) of the final model, shown in the same orientations as the top. Scale bars are 100 Å, unless stated otherwise. See Table S2 for image processing details. **a.** Orthogonal views of the HTL-SurA complex 3D reconstruction (41.9 Å resolution). **b.** Opposing side views of the HTL-SurA complex (yellow) superimposed on the immuno-complex HTL-SurA:Ab_SurA_ complex (grey). The antibody (yellow arrows) is shown bound to the mass assigned to SurA. **c.** Opposing side views of the HTL-SurA complex (yellow) superimposed on the immuno-complex Ab_SecY_:HTL-SurA complex (grey). The antibody (blue arrows) is shown bound to the mass assigned to SecYEG. **d.** Orthogonal side views of the HTL-SurA:OmpA complex 3D reconstruction (37.5 Å resolution). **e.** Assigned map of the HTL-SurA-OmpA complex in the three representative orthogonal views, with components coloured accordingly.

The experiment was repeated using SurA purified in complex with a classical OMP substrate, OmpA (Fig. 6d,e; Fig. S12a-b). The NS-EM structure of HTL-SurA-OmpA shows additional density compared to HTL-SurA, roughly adjacent to the channel exit site of SecYEG and in contact with SurA, possibly representing a late inner-membrane translocation intermediate (Fig. 7).

**Figure 7:**
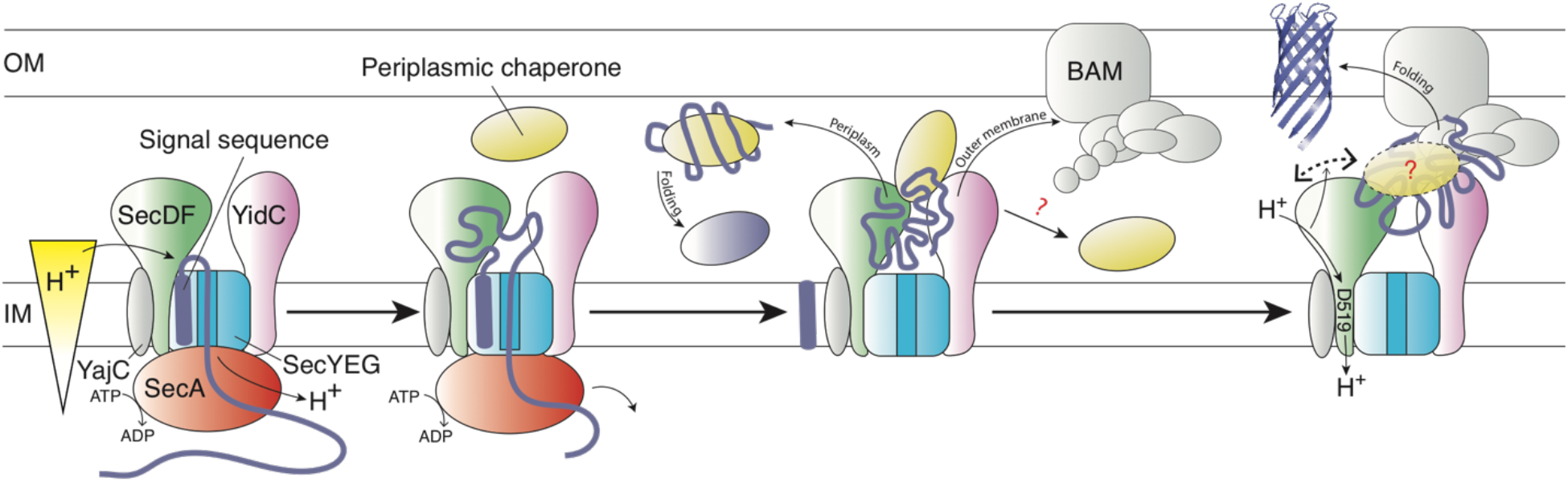
Schematic representation of the HTL:BAM machinery. Schematic model of OMP transfer through the bacterial envelope, facilitated by HTL-BAM and periplasmic chaperones, such as SurA, Skp, PpiD and YfgM. From left to right: OMP precursors with an N-terminal signal sequence are driven across the membrane by the ATPase SecA through the Sec translocon – the process is stimulated by PMF (independent of SecDF). Late in this process the pre-protein emerges into the periplasm and the signal sequence is removed, releasing the mature protein. Presumably, globular proteins are then guided into the periplasm, where folding will occur assisted by periplasmic chaperones. Otherwise, OMP-chaperone-HTL complexes are recognised by the BAM complex, with interactions forming between BAM and both HTL (this study) and SurA (Sklar et al., 2007). The persistence and variety of chaperones involvement at this stage is unclear (?). This conjunction enables the smooth and efficient passage of OMPs to the outer-membrane, enabled by energy coupling from the inner-membrane – trans-membrane proton-motive-force (PMF)-driven conformational changes *via* SecDF (right).

## DISCUSSION

The *in vivo* and *in vitro* analyses described here demonstrate a direct, functional interaction between the Sec and BAM translocons, mediated by the extended periplasmic domains possessed by BAM (Ma et al., 2019), SecDF (Furukawa et al., 2017) and YidC (Kumazaki et al., 2014), but not SecYEG. Evidently, direct contact between HTL and BAM is required for efficient OMP biogenesis in rapidly growing cells. The interaction could enable large protein fluxes to stream through the periplasm, while minimising aggregation and proteolysis (Fig. 7). The presence of a super-complex that bridges both membranes appears to be a fundamental feature of the Gram negative bacterial cell envelope, the importance of which is only just coming to the fore (Rassam et al., 2015; 2018).

It has already been shown that the HTL contains a lipid-containing cavity within the membrane, presumably to facilitate membrane protein insertion (Botte et al., 2016; Martin et al., 2019). Remarkably, in the super-complex between HTL and BAM there is a much larger extension of this cavity opening into the periplasm (Fig. 2). This would seem an obvious place for OMP passage and for the interaction with chaperones, and is of sufficient size to do so (Fig. 7). The cavity is situated such that secretory protein could enter *via* the protein-channel through SecYEG, and then exit accordingly into the periplasm or into the mouth of the BAM complex.

It remains to be seen how the Sec-BAM complex and the periplasmic chaperones coordinate. Perhaps these chaperones recognise emerging protein at the Sec-machinery and shuttle them into the periplasm, with or without the need for the BAM complex. Otherwise, they could facilitate the passage through the inter-membrane assembly for OM folding and insertion (Fig. 7). We show the interaction of SurA, as well as an OMP-SurA complex, with HTL; incriminating at least this chaperone with the early stages of transport through the cell wall, *i.e.* immediately after precursor cleavage. Furthermore, SurA is known to interact with BamA (Sklar et al., 2007), suggesting its involvement throughout OMP passage from the Sec to BAM (Fig. 7).

Other ancillary factors of the Sec machinery have also been implicated: YfgM and PpiD are thought to mediate interactions with periplasmic chaperones (Götzke et al., 2014), the latter of which has also been shown to interact with SecYEG and YidC (Jauß et al., 2019). Interestingly, *yfgL* and *yfgM* are in the same operon (Blattner et al., 1997); the former encoding a subunit of the BAM complex (BamB) (Wu et al., 2005). Indeed, a recent proteomic analysis of the *E. coli* ‘membrane protein interactome’ identifies cross-membrane interactions involving SecYEG, BAM and the chaperones YfgM and PpiD (Carlson et al., 2019). Understanding the interplay of various periplasmic chaperones during OMP passage through the Sec-BAM assembly to the outer membrane will require further attention.

From our data it is clear that the periplasmic domain of SecD is central to the physical BAM-HTL interaction. Even more intriguing though is the requirement for a functioning proton wire through the SecDF trans-membrane domain. This raises the intriguing prospect of a TonB-style energy-coupling from the inner membrane (Celia et al., 2016): *i.e.* the transmission of free energy available from ATP turnover and the PMF from the Sec-machinery (Arkowitz and Wickner, 1994; Brundage et al., 1990; Schiebel et al., 1991), for OMP folding and insertion at the outer-membrane. We therefore propose that one of the primary roles of SecDF is in inter-membrane trafficking and chemiosmotic energy transduction. Indeed, we and others have shown that ATP- and PMF-driven transport of proteins through the inner membrane is dependent only on SecYEG and SecA (Brundage et al., 1990; Schulze et al., 2014); while we show here proton translocation through SecD is crucial for efficient OMP folding and growth.

Thus, there appears to be two distinct requirements of the PMF in protein secretion: one for the early stage – SecA driven translocation through SecYEG at the inner membrane, and another for late stages of OMP maturation. The latter could be facilitated by conformational changes in SecDF, for transduction of energy from the inner to the outer membrane. When a key proton carrying residue of the inner membrane segment of the translocon is neutralised – SecD_D519N_ – the conformation of the periplasmic domain is perturbed (Furukawa et al., 2017). Presumably then, successive protonation (approximated by SecD_D519N_) and deprotonation result in large, long-range, cyclical conformational changes during PMF driven proton transport from the periplasm to the cytosol. Interestingly, the phospholipid cardiolipin (CL) is also important for the association of the Sec and BAM machineries; it is probably not a coincidence that this lipid has already been shown to be critical for PMF driven protein translocation through SecYEG (Corey et al., 2018).

Taking all together, this builds a compelling case for SecD mediated inter-membrane energy transduction for outer membrane protein folding and insertion – in keeping with other members of the RND transporter family, such as the assembly of AcrAB (inner membrane) and TolC (outer membrane) (Du et al., 2014; Wang et al., 2017). Direct association between inner and outer membrane components appears to be the rule rather than the exception for transporters embedded in double membrane systems: parallels with the translocation assembly module (TAM) for auto-transporter secretion (Selkrig et al., 2012) and the TIC-TOC import machinery of chloroplasts (Chen et al., 2018) are particularly striking, given the respective outer-membrane components (TamA and TOC75) are homologous of BamA. Particularly intriguing is the possibility of the mitochondrial homologue of BAM (Sorting and Assembly Machinery; SAM) participating in analogous inter-membrane interactions between respective inner and outer membranes. Indeed, subunits of the MItochondrial contact site and Cristae Organizing System (MICOS) connect the energy-transducing ATP synthase of the inner membrane and SAM at the outer-membrane (Ott et al., 2015; Rampelt et al., 2017).

## MATERIALS AND METHODS

### Strains and plasmids

*E. coli* C43 (DE3) was a gift from Sir John Walker (MRC Mitochondrial Biology Unit, Cambridge, UK) (Miroux and Walker, 1996). *E. coli* BL21 (DE3) were purchased as competent cells (New England Biolabs). *E. coli *Δ*secG* (KN425 (W3110 M25 *ΔsecG*::kan)) (Nishiyama et al., 1994), which lacks a genomic copy of *secG*, was obtained from Prof. Frank Duong (University of British Colombia, Vancouver, Canada)*. E. coli* strain JP352 (Kan^r^), which contains an arabinose-regulated *secDF-yajC* operon (Economou et al., 1995), was given to us by Prof. Ross Dalbey.

The plasmids for over-expression of *secEYG* and *yidC* were from our laboratory collection (Collinson et al., 2001; Lotz et al., 2008), the former and also that of *secDF* were acquired from Prof. Frank Duong (Duong and Wickner, 1997). Vectors designed for over-production of HTL, HTL(ΔYidC) and HTL(SecD_D519N_) were created using the ACEMBL expression system (Bieniossek et al., 2009; Botte et al., 2016). The vector for *bamABCDE* over-expression pJH114 (Amp^r^) was a gift from Prof. Harris Bernstein (Roman-Hernandez et al., 2014); from which pJH114-*bamACDE* (ΔBamB) was produced by linear PCR with primers designed to flank the BamB gene and amplify DNA around it. FseI restriction sites were included in the primers to ligate the amplified DNA. pBAD-SecDF(ΔP1) was generated by amplifying SecDF(ΔP1) from pBAD-SecDF and cloning it between the pBAD NcoI and HindIII sites. pET28b-*surA* and pET11a-*ompA* (Schiffrin et al., 2017) were gifts from Prof. Sheena Radford (The Astbury Centre for Structural Molecular Biology, University of Leeds, UK).

For SecDF depletion experiments, SecDF was cloned into pTrc99a (Amp^r^, IPTG-inducible), and the *secD*_*D519N*_ mutation was subsequently made by changing the WT carrying plasmid using a site-directed ligase-independent PCR method.

### SDS-PAGE, western blotting and antibodies

All SDS-PAGE was performed with either *Invitrogen* Bolt 4-12% Bis-Tris gels or *Invitrogen* midi 4-12% Bis-Tris gels. For western blotting, proteins were transferred onto nitrocellulose membrane. Mouse monoclonal antibodies against SecY, SecE and SecG were from our laboratory collection (used at 1:10000 dilution). Polyclonal antibodies against SecD and BamA were generated commercially in rabbits, and SecA in sheep from purified proteins (All used at 1:5000 dilution). BamB and BamD antibodies were gifts from Dr Harris Bertstein (1:5000 dilution). The SurA antibody was purchased from 2BScientific (1:10000 dilution). A secondary antibody conjugated to DyLight800 was used for SecG and SecY (Thermo Fisher Scientific, 1:10000 dilution), whereas a HRP-conjugated secondary antibody was used for SecD and BamA (1:10000 dilution).

### Protein production and purification

HTL, HTL(ΔYidC), HTL(SecD_D519N_), SecYEG, YidC, SecDF and SurA were purified as described previously (Burmann et al., 2013; Collinson et al., 2001; Lotz et al., 2008; Schulze et al., 2014). BAM and BAM(ΔBamB) was over-produced in *E. coli* C43 according to established protocols (Iadanza et al., 2016; Roman-Hernandez et al., 2014). For preparation of SurA and SurA-OmpA complexes, both proteins were over-produced separately in 1 L of cultures as described previously (Humes et al., 2019). Both were harvested by centrifugation, lysed in a cell disruptor (Constant Systems Ltd.) and resuspended in 20 mL 20 mM Tris pH 8.0, 130 mM NaCl, 10% (v/v) glycerol (TS_130_G) in the presence of cOmplete protein inhibitor cocktail (Roche). The resulting samples were clarified by centrifugation in an SS34 rotor (27,000 x*g*, 4°C, 20 minutes, Sorvall). For OmpA, the supernatant was discarded and the pellet resuspended in 20 mL TS_130_G + 6 M urea. The OmpA pellet was diluted to 80 mL with 6 M urea and mixed with the SurA supernant to give a final urea concentration of 4.8 M. The urea was removed by dialysing in 2 L TS_130_G for 6 hours at room temperature, then dialysing overnight at 4°C in 5 L fresh TS_130_G. The sample was centrifuged under the same conditions as above and the supernatant loaded onto a 5 mL HisTrap HP column equilibrated with TS_50_G. The column was washed with TS_50_G + 20 mM imidazole, and bound proteins eluted with TS_50_G + 300 mM imidazole. The eluents were loaded onto a HiTrap Q HP column equilibrated in TS_50_G and free SurA was found in the unbound fraction. A linear gradient of 0.05 - 1 M NaCl was applied over 60 mL and the fractions containing SurA-OmpA were taken for further analyses.

### Isolation of inner and outer membranes

1 L of *E. coli* cultures over-producing SecYEG, HTL, SecDF or SecDF(*Δ*P1) were produced as described previously (Collinson et al., 2001; Komar et al., 2016; Schulze et al., 2014). The harvested cell pellets were resuspended in 20 mL TS_130_G, homogenised with a potter, passed twice through a cell disruptor (Constant Systems Ltd.) for lysis and centrifuged to remove debris (SS34 rotor, Sorvall, 12,000 x*g*, 20 minutes, 4°C). The supernatant was taken and layered upon 20 mL TS_130_G + 20% (w/v) sucrose in a Ti45 tube and centrifuged (Ti45 rotor, Beckmann-Coulter, 167,000 x*g*, 120 minutes, 4°C). The pellet was taken, resuspended in 4 mL TS_130_G, homogenised with a potter and layered upon a sucrose gradient prepared in an SW32 centrifuge tube composed of 5 mL layers of TS_130_G + 55% (w/v), 48%, 41%, 34% and 28% sucrose. The sample was then fractionated by centrifugation (SW32 rotor, Beckmann-Coulter, 130,000 x*g*, 15 hours, 4°C). Upon completion, the light to heavy fractions were analysed by SDS PAGE and western blotting.

### Co-immunoprecipitations (co-IPs) with *E. coli* total membrane extracts

Membrane pellets of *E. coli* strains C43 (WT), C43 pJH114-*bamABCDE* (Amp^r^), *Δ*secG (Kan^r^), WT *lpp*, mutant *lpp_+21_* and JP325 (containing variants of pTrc as specified in text, cultures grown in glucose for depletion of endogenous SecDFyajC), were prepared as described previously (Collinson et al., 2001), with *bamABCDE* over-expression achieved as before (Roman-Hernandez et al., 2014). The pellets were resuspended in TS_130_G to 120 mg/mL, homogenised and solubilised with 0.5% DDM for 1 hour at 4°C. The solubilised material was clarified by ultra-centrifugation (160,000 x*g* for 45 mins) and the membrane extracts were analysed.

For co-IPs pulling on SecG antibody, 250 uL of protein G resin was washed in a spin column with 200 mM NaCl, 20 mM HEPES pH 8.0 (HS buffer), and blocked overnight in HS buffer + 2% (w/v) BSA at 4°C. Meanwhile, 7.5 μL of purified SecG monoclonal antibody was added to 500 μL of the membrane extracts and incubated overnight at 4°C. The following morning, the resin was washed thoroughly in HS buffer containing either 0.02% (w/v) DDM or 0.02% (w/v) DDM with 0.002% (w/v) CL, resuspended back to 250 μL and added to the 500 μL of membrane extract for three hours rotating gently at room temperature. The resin was separated from the extracts by centrifugation in a spin column at 2,000 x*g* for 1 minute, washed 7 times with 350 μL HS buffer, followed by one final wash with 150 μL HS buffer, which was collected in a fresh tube for analysis (to which 50 uL of 4x LDS sample buffer was added once collected). The bound material was then eluted by addition of 150 uL 1 x LDS sample buffer (to which an additional 50 uL of 1x LDS sample buffer was added once collected). Samples were analysed by SDS PAGE and western blotting.

For co-affinity adsorption by pulling on the hexa-histidine tag of recombinant BamA, 100 μL of nickel-charged chelating resin was added to 500 μL of membrane extracts and incubated for 5 minutes at room temperature. The resin was then separated from the extract and treated in the same way as described above but with TS_130_G + 0.02% (w/v) DDM/ 0.002% (w/v) CL + 30 mM imidazole (washing) or 300 mM imidazole (elution).

Statistical analyses were conducted using GraphPad Prism. An unpaired T-test was used to compare pull down samples (p value = 0.05, * = < 0.05, ** = < 0.01, *** = < 0.001, specific p values are stated in figure legends).

### *In vitro* assembly and purification of complexes for EM and XL-MS

All protein complexes visualized by negative stain EM were formed by incubating 5 μM of the respective proteins in binding buffer (20 mM HEPES pH 8.0, 250 mM NaCl, 0.03% (w/v) DDM/ 0.003% (w/v) CL) at 30°C for 30 minutes with shaking in a total volume of 150 μl. The protein complexes were purified in a glycerol/ glutaraldehyde gradient (20 - 40% (w/v) and 0 - 0.15% (w/v), respectively) by centrifugation at 34,000 RPM in a SW60 Ti rotor (Beckmann-Coulter) for 16 hours at 4°C. Mobility controls of individual and partial complexes (BAM, and HTL) or individual proteins (SecYEG, YidC, SecDF, SurA) without the glutaraldehyde gradient were performed under the same conditions. Gradients were fractionated in 150 μl aliquots and those with glutaraldehyde were inactivated with 50 mM of Tris pH 8.0. Aliquots were analysed by SDS-PAGE and silver staining. For HTL:SurA:Ab_SurA_ and Ab_SecY_:HTL:SurA complexes, 5 μM of HTL and SurA were incubated in binding buffer at 30°C for 30 minutes with shaking in a total volume of 150 μl. Antibodies were added at concentrations of 1 in 7.5 or 1 in 15 for the SecY or SurA antibodies respectively, these were incubated at 30°C for 15 minutes before complex purification as described above.

The HTL:BAM complex for cryoEM was formed by incubating 8 μM of the HTL and BAM complexes in binding buffer (50 mM HEPES pH 8.0, 200 mM NaCl, 0.01% (w/v) DDM / 0.001% (w/v) CL at 30°C for 20 minutes with shaking in a total volume of 250 μl. After 20 min, 0.05% of glutaraldehyde was added to the sample and incubated for 10 minutes at 21°C. The crosslinker was inactivated with 30 mM Tris pH 8.0 and the sample was loaded onto a Superose 6 Increase 10/300 GL (GE healthcare) column equilibrated in GF buffer (30 mM Tris pH 8.0, 200 mM NaCl, 0.01% (w/v) DDM). Fractions were analysed by SDS-PAGE and silver staining.

The HTL:BAM complex for cross-linked mass spectroscopy (XL-MS) analysis was prepared following the same procedure described for the cryo-EM preparation, but the sample was crosslinked with 1.5 mM DSBU and inactivated with 20 mM of ammonium carbonate pH 8.0 before loaded onto the gel filtration column.

### XL-MS analysis

The DSBU cross-linked HTL:BAM complex was precipitated by methanol and chloroform (Wessel and Flügge, 1984) and the pellet dissolved in 8 M urea. After reduction with 10 mM DTT (one hour at 37°C) and alkylation with 50 mM iodoacetamide (30 minutes in the dark at RT), the sample was diluted 1:5 with 62.5 mM ammonium hydrogen carbonate and digested with trypsin (1:20 w/w) overnight at 37°C. Digestion was stopped by the addition of formic acid to a final concentration of 2% (v/v) and the sample split in two equal amounts for fractionation by size exclusion (SEC) and reverse phase C18 at high pH chromatography. A Superdex Peptide 3.2/300 column (GE Healthcare) was used for SEC fractionation by isocratic elution with 30% (v/v) acetonitrile/ 0.1% (v/v) TFA at a flow rate of 50 μL/ min. Fractions were collected every minute from 1.0 mL to 1.7 mL of elution volume. Reverse phase C18 high pH fractionation was carried out on an Acquity UPLC CSH C18 1.7 μm, 1.0 x 100 mm column (Waters) over a gradient of acetonitrile 2-40% (v/v) and ammonium hydrogen bicarbonate 100 mM.

All the fractions were lyophilized and resuspended in 2% (v/v) acetonitrile and 2% (v/v) formic acid for LC–MS/MS analysis. An Ultimate U3000 HPLC (ThermoScientific Dionex, USA) was used to deliver a flow of approximately 300 nL/ min. A C18 Acclaim PepMap100 5 μm, 100 μm × 20 mm nanoViper (ThermoScientific Dionex, USA), trapped the peptides before separation on a C18 Acclaim PepMap100 3 μm, 75 μm × 250 mm nanoViper (ThermoScientific Dionex, USA). Peptides were eluted with a gradient of acetonitrile. The analytical column was directly interfaced *via* a nano-flow electrospray ionisation source, with a hybrid quadrupole orbitrap mass spectrometer (Q-Exactive HF-X, ThermoScientific, USA). MS data were acquired in data-dependent mode. High-resolution full scans (R=120,000, m/z 350-2000) were recorded in the Orbitrap and after CID activation (stepped collision energy 30 ± 3) of the 10 most intense MS peaks, MS/ MS scans (R=45,000) were acquired.

For data analysis, Xcalibur raw files were converted into the MGF format through MSConvert (Proteowizard (Kessner et al., 2008)) and used directly as input files for MeroX (Götze et al., 2015). Searches were performed against an ad-hoc protein database containing the sequences of the complexes and a set of randomised decoy sequences generated by the software. The following parameters were set for the searches: maximum number of missed cleavages 3; targeted residues K; minimum peptide length 5 amino acids; variable modifications: carbamidomethyl-Cys (mass shift 57.02146 Da), Met-oxidation (mass shift 15.99491 Da); DSBU modification fragments: 85.05276 Da and 111.03203 (precision: 5 ppm MS (Kessner et al., 2008) and 10 ppm MS (Götze et al., 2015)); False Discovery Rate cut-off: 5%. Finally, each fragmentation spectra were manually inspected and validated.

### EM and image processing

For negative stain, aliquots of sucrose gradient fractions containing the different complexes were applied to glow-discharged (15 s) carbon grids with Cu 300 mesh, washed and stained with 2% (w/ v) uranyl acetate (1 min). Digital images were acquired with two different microscopes; a Tecnai 12 with a Ceta 16M camera (ThermoFisher Scientific) at a digital magnification of 49,000 x and a sampling resolution of 2.04 Å per pixel, and in a Tecnai 12 with a Gatan Camera One View at a digital magnification of 59,400 and a sampling resolution of 2.1 Å per pixel. Image processing was performed using the EM software framework Scipion v1.2 (la Rosa-Trevín et al., 2016). Several thousand particles were manually and semi-automatic supervised selected as input for automatic particle picking through the XMIPP3 package (Abrishami et al., 2013; la Rosa-Trevín et al., 2013). Particles were then extracted with the Relion v2.1 package (Kimanius et al., 2016; Scheres, 2012a) and classified with a free-pattern maximum-likelihood method (Relion 2D-classification). After manually removing low quality 2D classes, a second round of 2D classification was performed with Relion and XMIPP-CL2D in parallel (Sorzano et al., 2010). Representative 2D averages were used to generate several initial 3D models with the EMAN v2.12 software (Scheres, 2012b; Tang et al., 2007). Extensive rounds of 3D classification were then carried out using Relion 3D-classification due to the heterogeneity of the sample. The most consistent models were used for subsequent 3D classifications. For the final 3D volume refinement, Relion auto-refine or XMIPP3-Projection Matching were used. Resolution was estimated with Fourier shell correlation using 0.143 correlation coefficient criteria (Rosenthal and Henderson, 2003; Scheres and Chen, 2012). See Table S2 for image processing details.

For CryoEM, appropriate fractions of the glutaraldehyde-crosslinked HTL:BAM complex purified by gel filtration were applied to glow-discharged (20 s) Quantifoil grids (R1.2/R1.3, Cu 300 mesh) with an ultrathin carbon layer (2 nm), blotted and plunged into a liquid ethane chamber in a Leica EM. Two data sets from the same grid were acquired in a FEI Talos Arctica cryo-electron microscope operated at 200 kV and equipped with a K2 detector at calibrated magnification of 79000 x. The first data set with 2056 images recorded, had a 1.75 Å/px sample resolution, dose rate of 2.26 electrons/Å^2^ and 20 s exposure time fractionated in 40 frames. Defocus values oscillated between −1.5 nm and −3.0 nm. The second data set with 3703 images recorded, had a 0.875 Å/px sample resolution, dose rate of 2.47 electrons/Å^2^ and 18 s exposure time fractionated in 40 frames. Defocus values oscillated between – 1.0 nm and −2.2 nm. Particles were picked in the same way as for negative stain, and were binned to a 1.75 Å/px sample resolution before merging to the first data set. Image processing was performed using the EM software framework Scipion v1.2 (la Rosa-Trevín et al., 2016) with a similar strategy of the NS-EM samples but also using extensive masking procedures (Fig. S6).

All 3D reconstructions were calculated using a home-built workstation (CPU Intel Core i7 7820X, 2x Asus Turbo GTX 1080Ti, 16Gb RAM DDR4) and partial usage of HPC clusters (Bluecrystal 4 and Bluecryo) at the University of Bristol.

### Depletion of SecDF-YajC

*E. coli* strain JP352 was transformed with an empty pTrc99a, or the same plasmid, but cloned with either WT *secDF* or s*ecD*_*D519N*_*F*. Precultures of the strains were prepared in 100 mL 2xYT media supplemented with 0.2% (w/v) arabinose, ampicillin (for pTrc selectivity) and kanamycin (for JP352 selectivity). The following morning, the cells were harvested by centrifugation and resuspended with 50 mL fresh 2xYT (no arabinose). This washing procedure was repeated two more times to remove excess arabinose. Prewarmed (37°C) 1 L 2xYT cultures containing either 0.2% (w/v) arabinose or 0.5% (w/v) glucose were then inoculated with the preculture such that a final OD_600 nm_ of 0.05 was achieved. An aliquot was taken every 1.5 hours for 6 hours. Induction of pTrc with IPTG was not necessary as background expression was sufficient to achieve levels of SecD similar to that of JP325 cultured in the presence of arabinose. Periplasmic and outer membrane fractions were produced by preparing spheroplasts (Birdsell and Cota-Robles, 1967), centrifuging the samples at 12000 x*g* for 5 minutes, taking the supernatant (a mixture of periplasmic and OM fractions) and removing the OM fraction by ultracentrifugation at 160000 x*g* for 20 minutes. The fractions were then subjected to SDS-PAGE and western blotting.

### Measurement of protein transport

Inner membrane vesicles (IMVs) were produced from BL21(DE3) cells overproducing HTL, HTL(SecD_D519N_), SecYEG or with empty pBAD as described previously (Corey et al., 2018). Transport experiments with and without PMF were performed in triplicate using established methods (Corey et al., 2018).

**Table 1:**
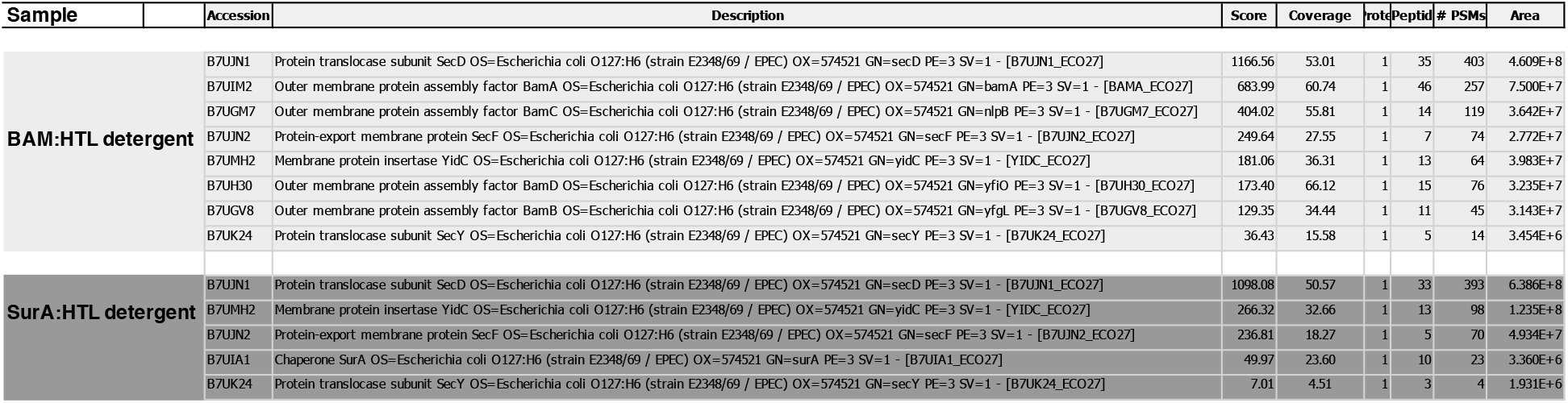
Mass spectrometry analysis of the GraFix fractions for image processing. Analysis of the protein composition of the HTL-BAM and HTL-SurA preparations in detergent solution by MS. Only proteins of interest are shown in this table. The SDS-PAGE bands corresponding to the GraFix fractions used for NS-EM processing (*e.g.* Fig. S4a, middle and right, asterisk) were cut from the gel and prepared for protein digestion, extraction and MS.

**Table 2:** Parameters of EM analysis of HTL, HTL-BAM and HTL-SurA structures.

|  | Complex | Initial particles | High quality particles | Final 3D volume | Resolution |
| --- | --- | --- | --- | --- | --- |
| HTL structures | HTL-DDM | 100754 | 74070 | - | - |
|  | HTL-DDM+CL | 22869 | 18674 | - | - |
|  | HTLafter gradient+CL | 21735 | 18434 | - | - |
|  | HTLafter gradient+gluta.+CL | 101142 | 90432 | 9656 (compact conformation) | 26,56 Å |
|  |  |  |  | 1569 (extended conformation) | 39,54 Å |
|  | HTL519<sub>D519N</sub> | 42098 | 38052 | 20699 (compact conformation) | 18,7 Å |
|  |  |  |  | 4451 (extended conformation) | 20,70 Å |
| HTL-BAM structures | HTL:BAM Δgluta | 29336 | 12460 | - | - |
|  | HTL:BAM grafix | 65381 | 28646 | 5122 | 37,2 Å |
| | HTL:BAM grafix ΔCL | 14530 | 11406 | - | - |
|  | HTL:BAM CryoEM | 839845 | 608493 | 12779 | 18,23 Å |
|  | SecDF:BAM | 22642 | 11987 | 5430 | 39,4 Å |
| | HTLΔYidC:BAM | 16008 | 8612 | 4943 | 36,7 Å |
| | HTL:BAMΔB | 31561 | 11484 | 3919 | 33,65 Å |
| HTL-SurA structures | SurA:HTL | 24252 | 18696 | 2290 | 41,88 Å |
|  | SurA:HTL:Ab<sub>SecY</sub> | 15819 | 9510 | 4431 | 47,06 Å |
|  | Ab<sub>SurA</sub>:SurA:HTL | 14246 | 3882 | 730 | 56,67 Å |
|  | SurA:HTL:OmpA | 35546 | 33485 | 11699 | 37,50 Å |

## Acknowledgments

We are particularly grateful for the generosity of Dr Harris Bernstein for the kind gifts of the *bamABCDE* expression construct (pJH114) and antibodies, and Prof. Sheena Radford for *surA* and *ompA* expression plasmids. Thanks to Prof. Daniel Daley for telling us about YfgL and YfgM. We thank Dr Remy Martin for brewing, and for making the HTL *SecD_D519NF_* mutant. We acknowledge access and support of the Wolfson Bioimaging Facility and the GW4 Facility for High-Resolution Electron Cryo-Microscopy, with particular thanks to Dr Ufuk Borucu. We are grateful to Mr Tom Batstone for the support at the computer cluster BlueCryo.

## Author contribution

SA, DWW, LT, WA and IC designed and conducted experiments, assisted by JL; SA, DWW, LT and IC wrote the manuscript; VAMG and BD provided facilities and expertise, and edited the manuscript; EJC and MB, helped with the Lpp experiments – conception, strain provision and text editing; GD and JMS, conducted the XL-MS experiments, processed the data, helped in their description and interpretation; IC, VAMG, BD and MB secured funding; IC led the project.

## Declaration

the authors declare no competing interests.

## Data and materials availability

All data are available in the main text or the supplementary materials. The HTL-BAM cryo-EM structure has been deposited at the EMDB under the accession number 11240.

---

**Supplementary figure 1:**
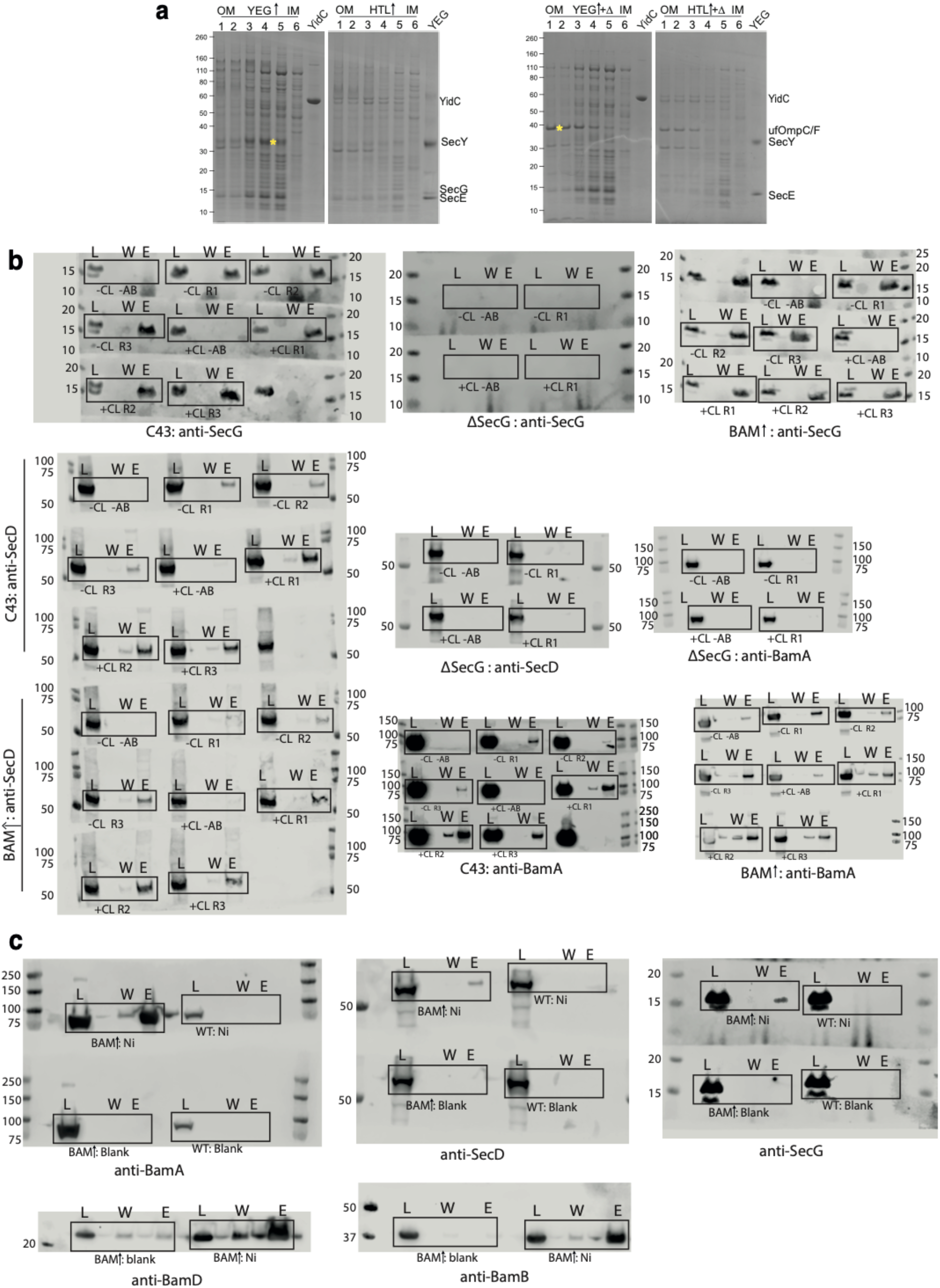
Raw western blots of co-immuno-precipitations and affinity pull downs. **a.** SDS PAGE stained for total protein of sucrose gradient fractions of *E. coli* C43 total membranes from cells over-producing SecYEG or HTL. Left and right gels show untreated and heat denatured (+Δ) samples respectively. Purified controls of SecYEG and YidC are also shown. For (**b)** and **(c)**, L = load (1% total material), W = final wash before elution (to demonstrate complete washing of affinity resin, 17% of total material) and E = elution (17% of total material). **b.** Co-immunoprecipitations from Fig. 1d-e. Experiments were conducted in the presence (+CL) or absence (-CL) of augmented cardiolipin. Experiments were also carried out in the presence (+AB) or absence (-AB) of anti-SecG primary antibody, as a control to assess non-specific binding. Solubilised membranes of 3 cell strains were used: *E. coli* C43, a strain lacking SecG (*ΔsecG*), and C43 over-producing BAM. **c.** Affinity pull-downs from Fig. 1f. Experiments were conducted in with nickel-chelated (Ni) or non-chelated (blank) resin as a control to rule out non-specific binding.

**Supplementary figure 2:**
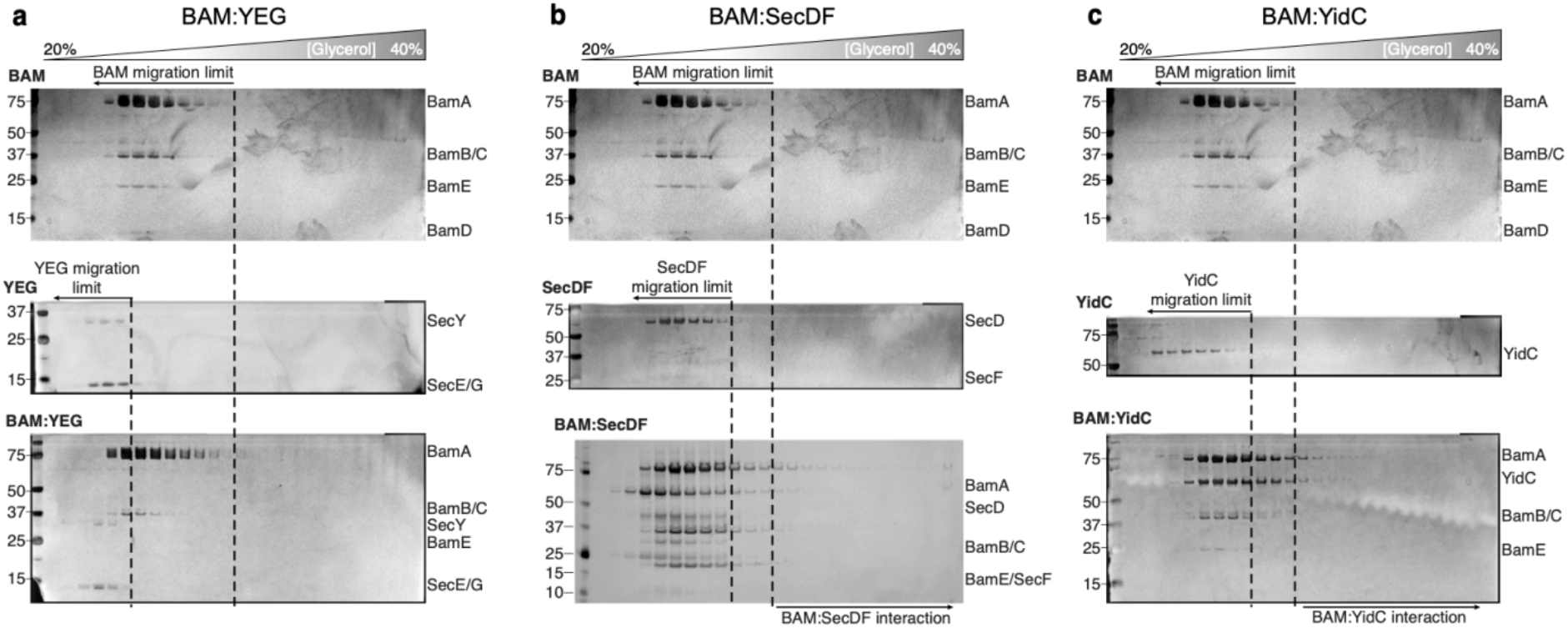
Glycerol centrifugation gradients of HTL and BAM components. **a, b, c.** Silver-stained SDS-PAGE gels of fractions from the glycerol centrifugation gradients are shown, with increasingly large complexes appearing in fractions of higher percentage glycerol, (from left to right). For each panel, the glycerol centrifugation gradient of BAM alone is shown (top). The middle gel represents a HTL component (labelled on the top left of the gel). The bottom gel represents the experiment where BAM was mixed with the corresponding HTL component. Dashed lines represent the fraction of furthest migration of the individual components, as determined in the top two gels.

**Supplementary figure 3:**
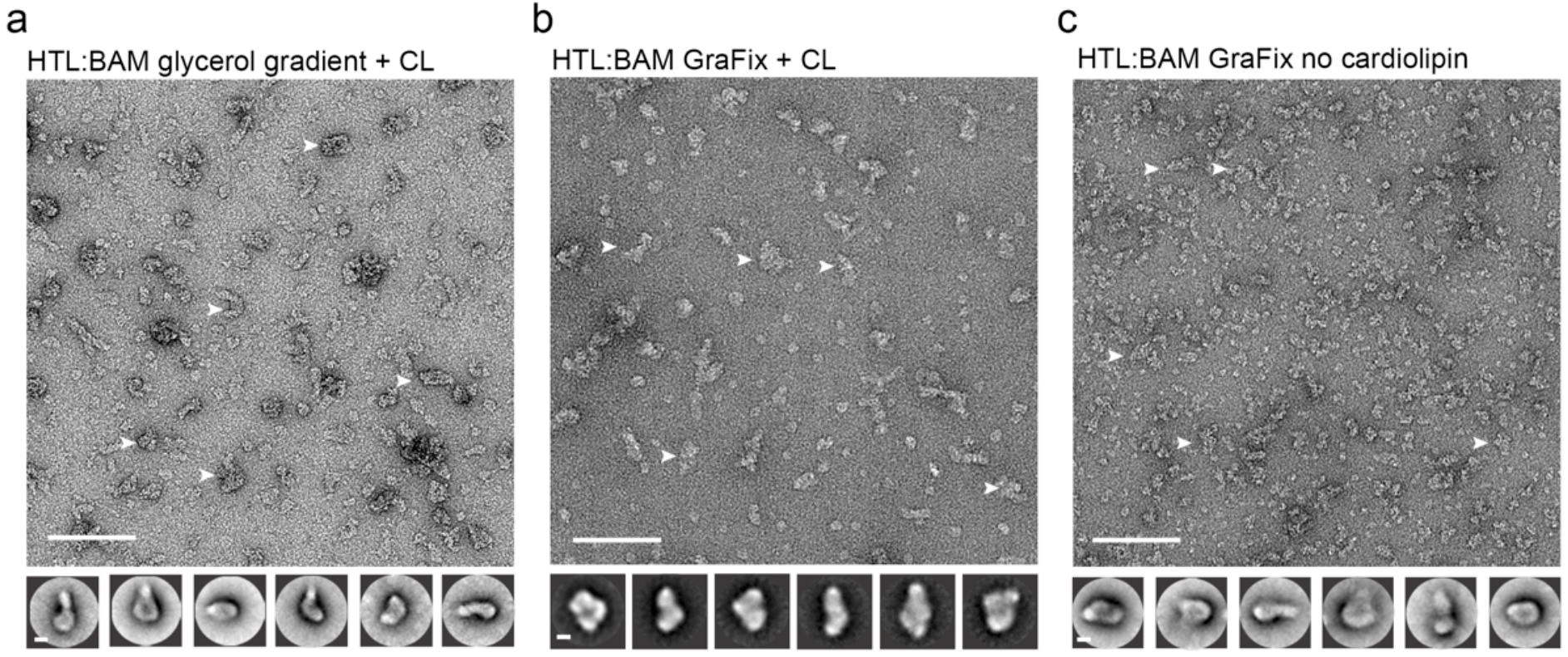
Negative-stained EM micrographs of the HTL:BAM complex. **a, b, c.** Electron micrographs of HTL-BAM complexes in different conditions. Bottom, reference-free (RF) class averages of the largest populations found in the micrographs (top). Micrograph scale bar, 1000 Å. RF scale bar, 100 Å. White arrows indicate representative HTL:BAM complexes used for image processing.

**Supplementary figure 4:**
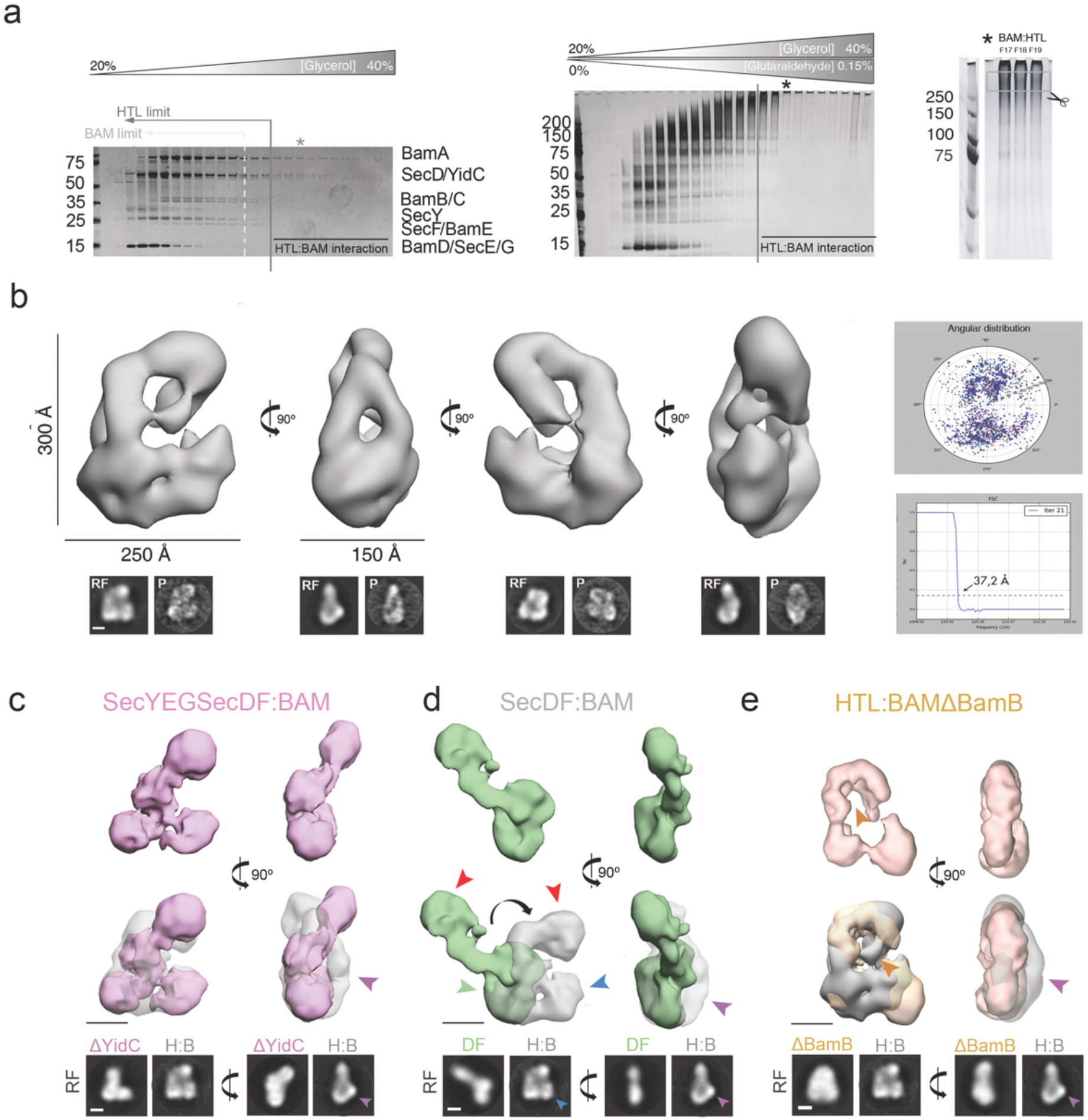
3D characterization and subunit assignment of HTL-BAM by NS-EM in detergent solution. **a.** Silver stained SDS-PAGE analysis of HTL-BAM fractionated by glycerol density gradient centrifugation (left), and the same but with GraFix treatment (middle). Lines show the limit to which BAM (dashed white line) and HTL (solid black line) migrate when ran alone (Fig. 2a). Asterisk (middle panel) shows the fraction chosen for MS (Table S1), EM and image processing. Right panel, silver stain SDS-PAGE gel showing fractions and bands used for analysis by mass spectrometry. **b.** Top left, four orthogonal views of the HTL-BAM complex 3D reconstruction (37.2 Å resolution). Bottom, reference-free (RF) class averages and projections (P) of the final model, shown in the same orientations as the top. Scale bar, 100 Å. See Table S2 for image processing details. Right, angular distribution and Fourier shell correlation of the HTL-BAM complex. For **(c-e)**, top panel shows orthogonal views of different 3D reconstructions, labelled accordingly. Middle panel shows reconstructions superimposed with HTL-BAM (transparent grey) from (**b),** and bottom shows a comparison of the reference-free (RF) class averages of HTL-BAM with the corresponding structure. Scale bar, 100 Å. See Table S2 for image processing details. **c.** SecYEG-SecDF-BAM (without YidC) complex 3D reconstruction (36.7 Å resolution, pink). Pink arrow indicates the mass in HTL-BAM (grey transparent) corresponding to YidC. **d.** SecDF-BAM (without SecYEG and YidC) complex 3D reconstruction (39.4 Å resolution, green). Masses for SecDF trans-membrane domain (green arrow) and periplasmic region (red arrow), SecYEG (blue arrow) and YidC (pink arrow) are indicated. **e.** HTL-BAM(ΔBamB) complex 3D reconstruction (33.6 Å resolution, orange). Mass corresponding to BamB (orange arrow) and YidC (pink arrow) are indicated.

**Supplementary figure 5:**
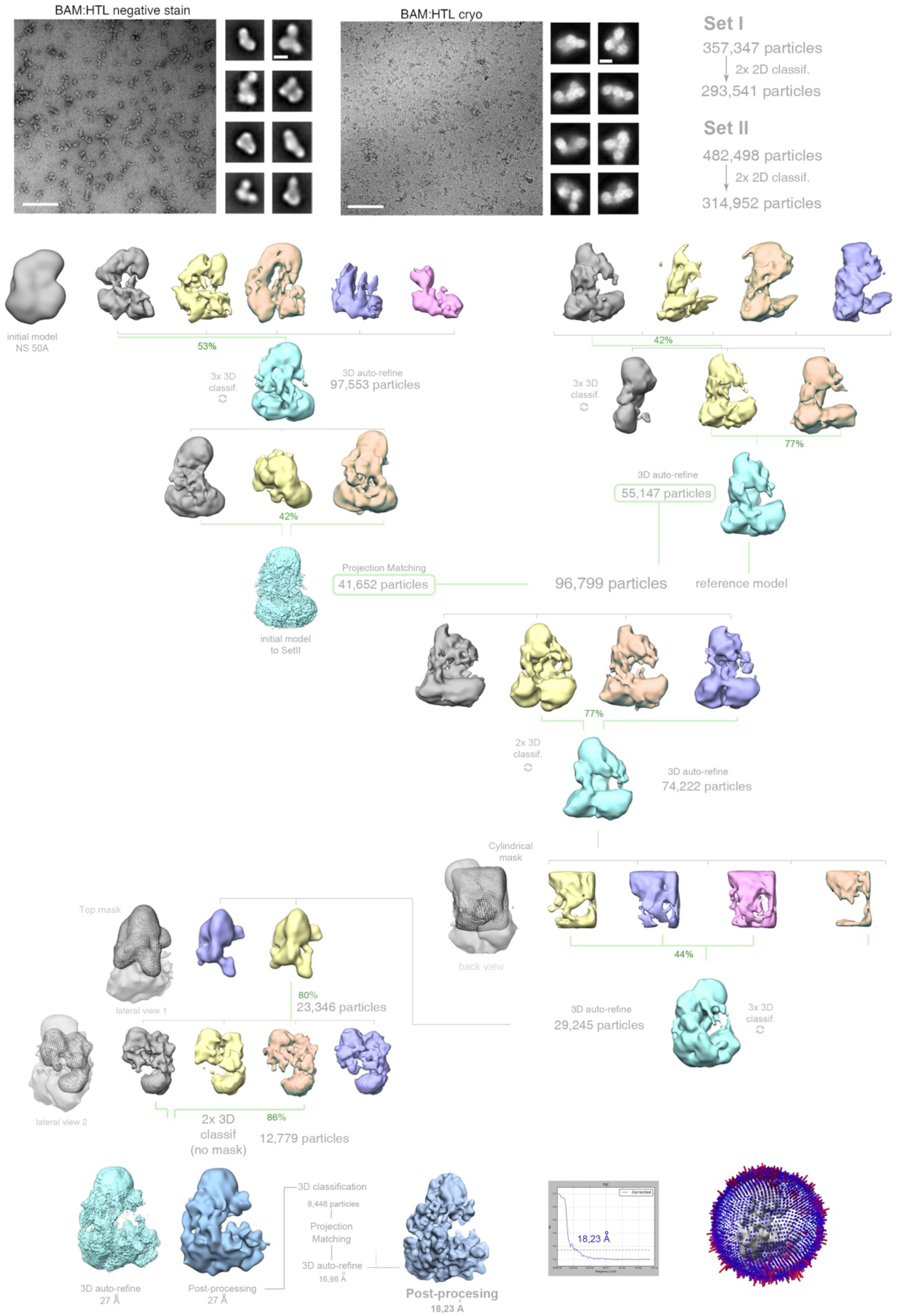
Image processing and classification strategy for the cryo-EM data of the HTL-BAM complex. Relion 2D classification was used to clean the two preliminary datasets and classify the images. Of Set I, 82% of the particles were used for extensive 3D classifications and auto-refinements. Of Set II, 65% of the particles followed same procedure. High-quality particles from both data sets were grouped and classified again masking the most variable regions. The final stable volume was formed by 8,448 particles (1.3% of the particles): indicative of the number of different sub-populations found in the sample. The final post-processing structure (Relion 2.0) has an overall resolution of 18,23 Å.

**Supplementary figure 6:**
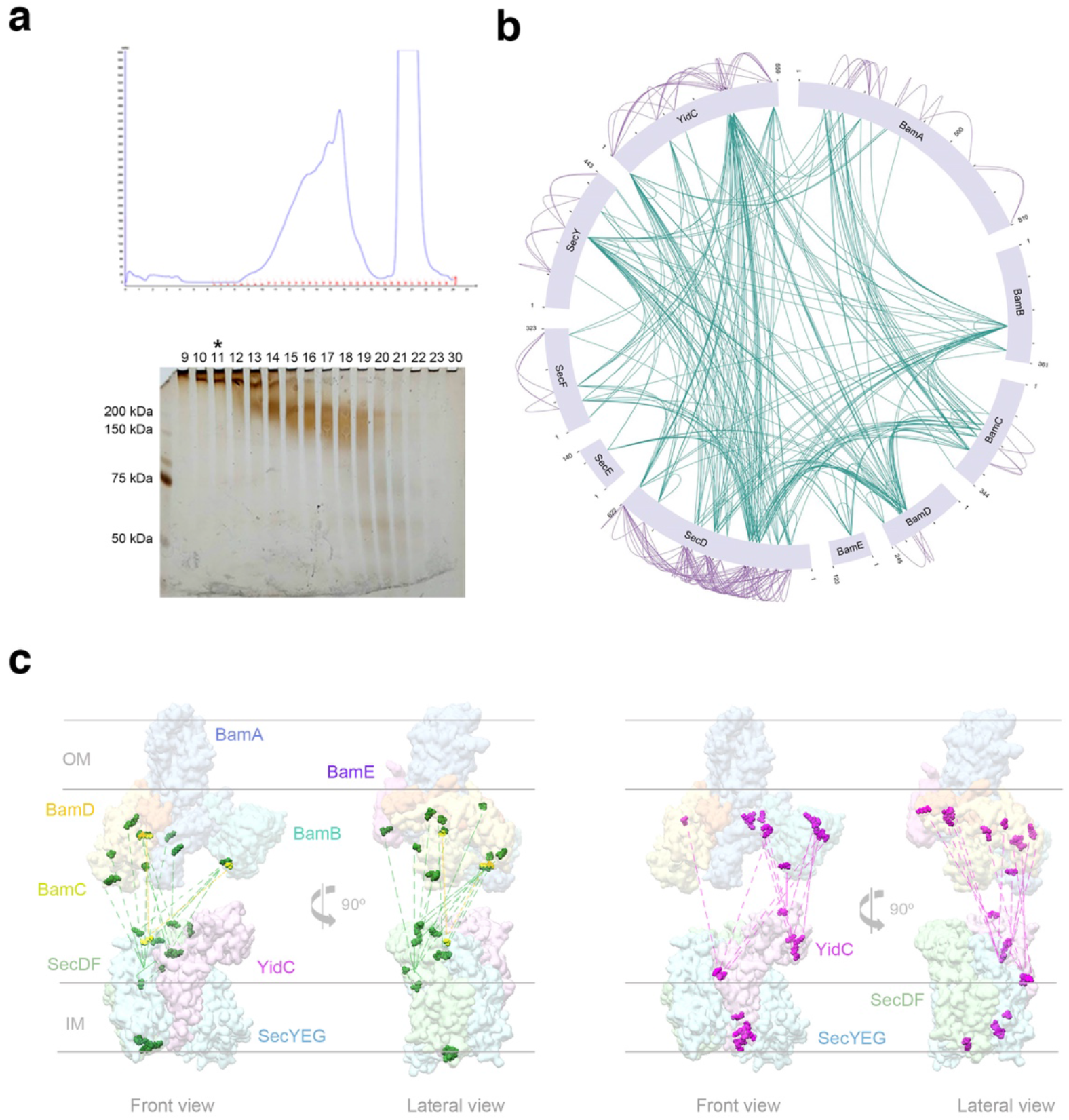
Sample preparation and XL-MS analysis of HTL-BAM. **a.** Top, Gel filtration chromatography profile elution (Superose 6) of the HTL-BAM complex for cryo-EM and XL-MS analysis. Bottom, SDS-PAGE gel silver stained of the fractions eluted from the gel filtration at the top. **b.** Representation of the interactome network between HTL and BAM resulted from XL-MS analysis. Internal green lines represent inter-connections between HTL-HTL, HTL-BAM and BAM-BAM components. Purple lines represent intra-connections between the HTL and BAM individual subunits. **c.** Representation of inter-protein interactions between HTL and BAM as determined by XL-MS. The specific crosslinks residues are mapped onto the structures of the BAM complex (emd_4061 and pdb_5ayw) and the model of the HTL complex (emd_3506 and pdb_5mg3) (Botte et al., 2016). Left, green dots and dashed lines represent positions and interactions, respectively, between SecD residues and BAM components. Yellow, positions and interactions between SecF and BAM components. Right, pink dots and dashed lines represent positions and interactions, respectively, between YidC residues and BAM components. The HTL and BAM structures have been placed artificially separate from one another to best illustrate the cross-linking contacts (rather than to reflect their actual position in the super-complex).

**Supplementary figure 7:**
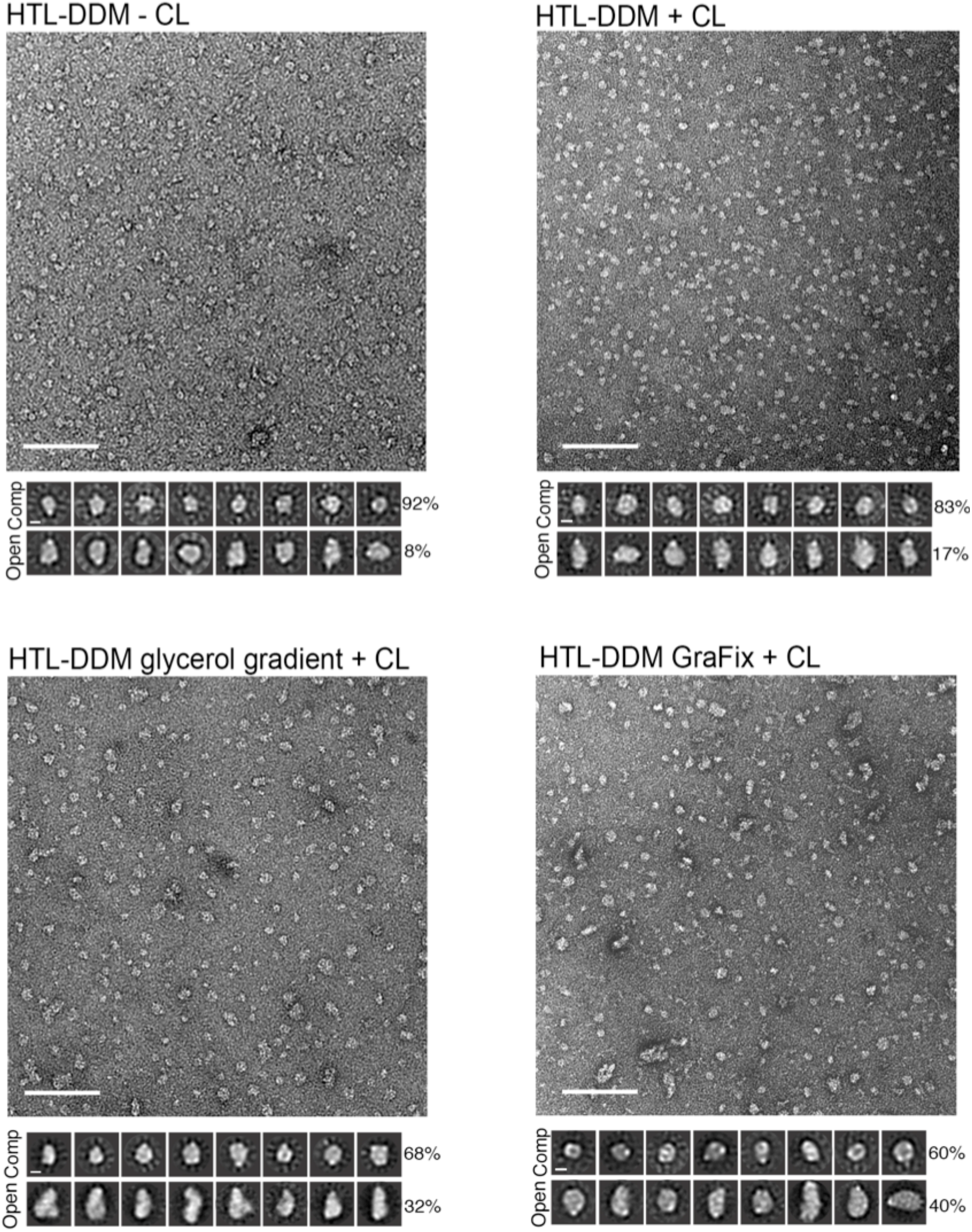
EM field of wild-type HTL in different conditions. Bottom, reference-free (RF) class averages of the ‘compact’ (comp) and ‘open’ populations found in the micrographs; conditions as labelled. Percentages of the populations are indicated on the right RF images. Micrograph scale bar, 1000 Å. RF scale bar, 100 Å.

**Supplementary figure 8:**
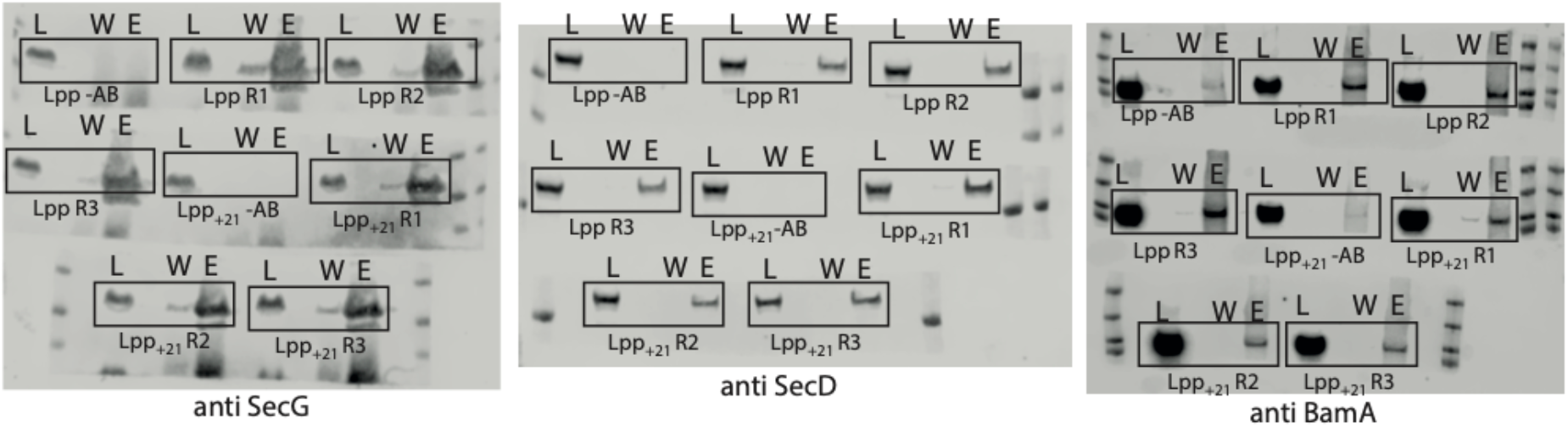
Raw western blots of IPs investigating how periplasmic width effects the HTL-BAM interaction. Legend is the same as described for Fig. S1a, but the cell strains used here were either *E. coli* containing WT *lpp* or the mutant *lpp_+21_*. All IPs were conducted in the presence of cardiolipin.

**Supplementary figure 9:**
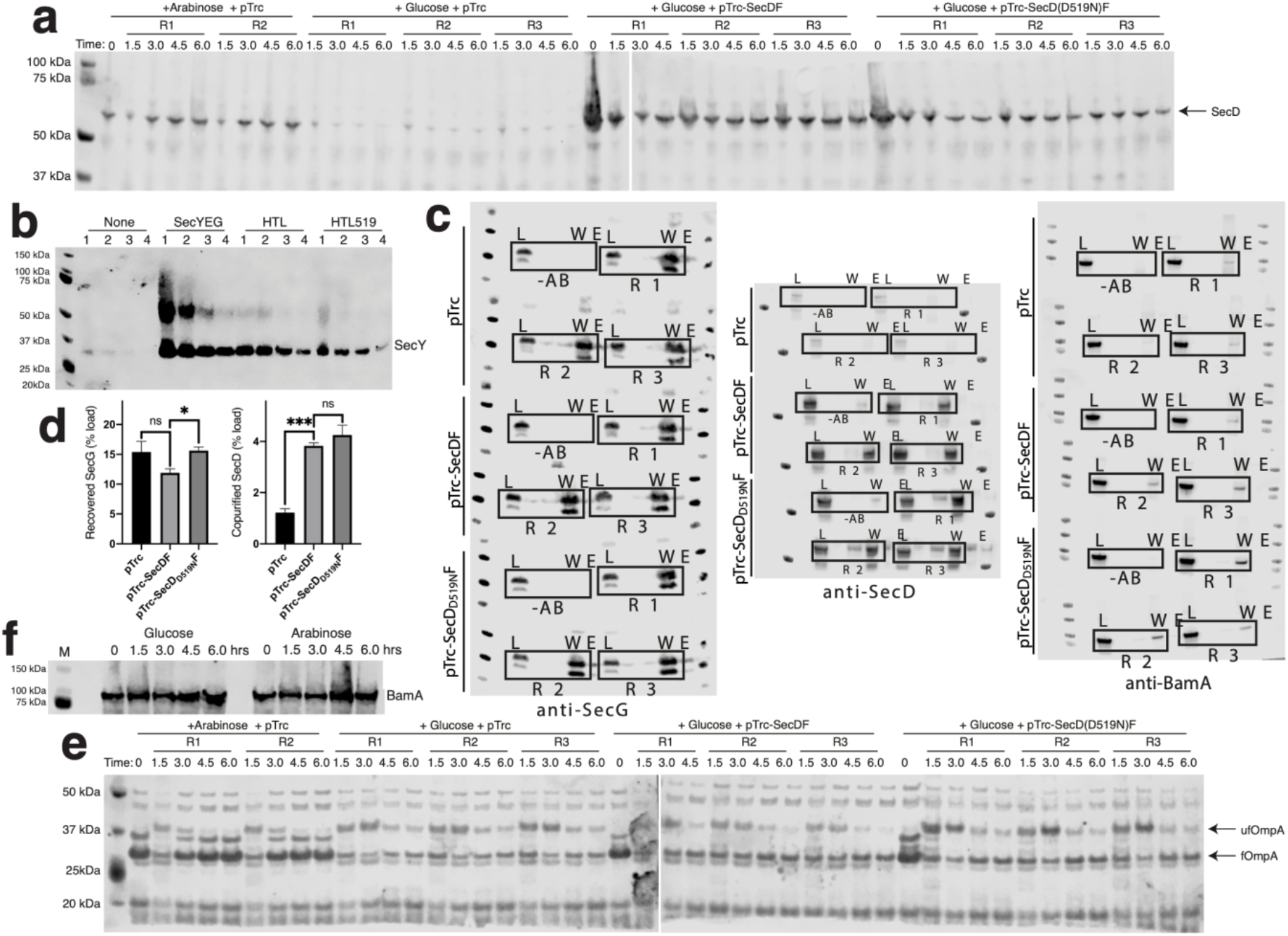
Raw western blots accompanying SecD depletion experiments from Fig. 5. **a.** Western blots of whole cells visualised for SecD. Cell strains used were *E. coli* JP325 containing either empty pTrc99a, pTrc99a-SecDF or pTrc99a-SecD_D519N_F. Samples were grown in either arabinose (permissive) or glucose (non-permissive), as indicated in the figure. R1/2/3 represent samples taken from replicate cultures. These blots were quantified to give Fig. 5f. **b.** Immunoblotting of SecY in inverted membrane vesicles used in Fig. 5c. Samples were prepared at 4 dilutions: neat (1), ½ (2), ¼ (3) and 1/8 (4). SecY is indicated at the expected apparent Mw of 30 kDa. A higher band is also present at approximately 50 kDa, likely due to SecY dimers. **c.** Raw western blots of co-IPs conducted in Fig. 5e-f. Legend is the same as for Fig. S1a, except the membranes used here were the same as those described in **(a**), all grown in the presence of glucose for depletion of genome encoded SecDF-yajC. All IPs were conducted in the presence of cardiolipin. **d.** Quantification of SecG and SecD from co-IPs shown in **(c)**. An unpaired t-test was used to compare samples (p = 0.05, n = 3, * = < 0.05, *** = < 0.001, p values from left to right are 0.0.1391, 0.0137, 0.0001 and 0.3618). **e.** OmpA western blots of periplasmic fractions of cells from **(a).** Unfolded (ufOmpA) and folded OmpA (fOmpA) are indicated. These blots were quantified to give Fig. 5i. **f.** BamA western blot of whole cells taken from cultures grown during depletion of SecDF-yajC.

**Supplementary figure 10:**
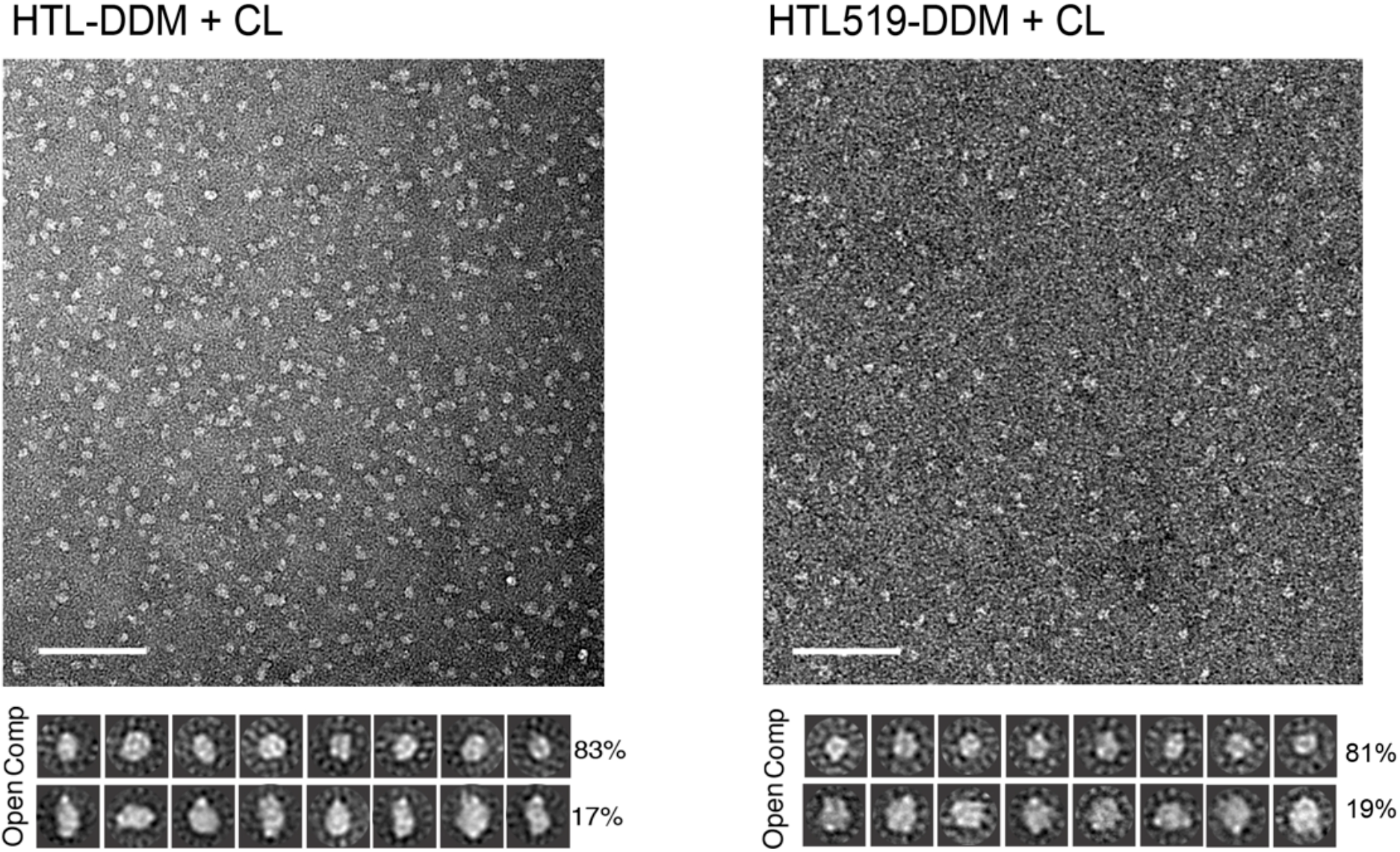
Negative-stain EM of HTL containing SecDD519N (HTL519) Electron micrographs of wild type (wt) HTL and HTL containing SecD_D519N_F (HTL519) in the same conditions (with cardiolipin, and without GraFix). Bottom, reference-free (RF) class averages of the ‘compact’ (comp) and ‘open’ populations found in the micrographs (top). Percentages of the populations are indicated on the right of RF images. Micrograph scale bar, 1000 Å. RF scale bar, 100 Å.

**Supplementary figure 11:**
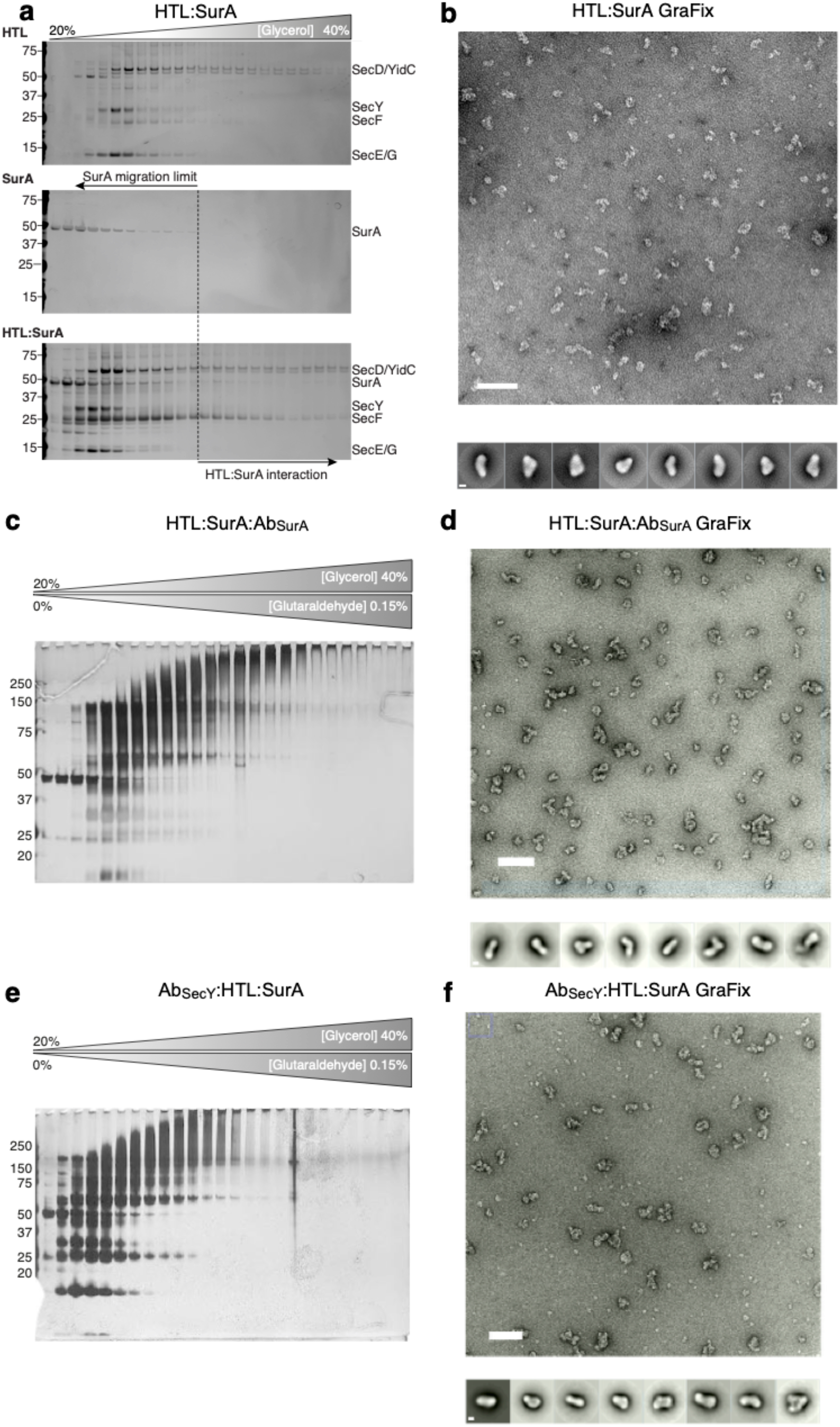
Glycerol gradient centrifugation, electron micrographs and immunodecoration of HTL:SurA. For **(a), (c)** and **(e)**, the experiment was conducted as in Fig. S2. EM fields of HTL:SurA (**b,** top) and HTL:SuraA:Ab_SurA_ (**d,** top) and Ab_SecY_:HTL:SurA (**f**, top) prepared using GraFix are shown. Reference-free (RF) class averages of the biggest populations from corresponding micrographs are shown (bottom). Micrograph scale bar, 1000 Å. RF scale bar, 100 Å.

**Supplementary figure 12:**
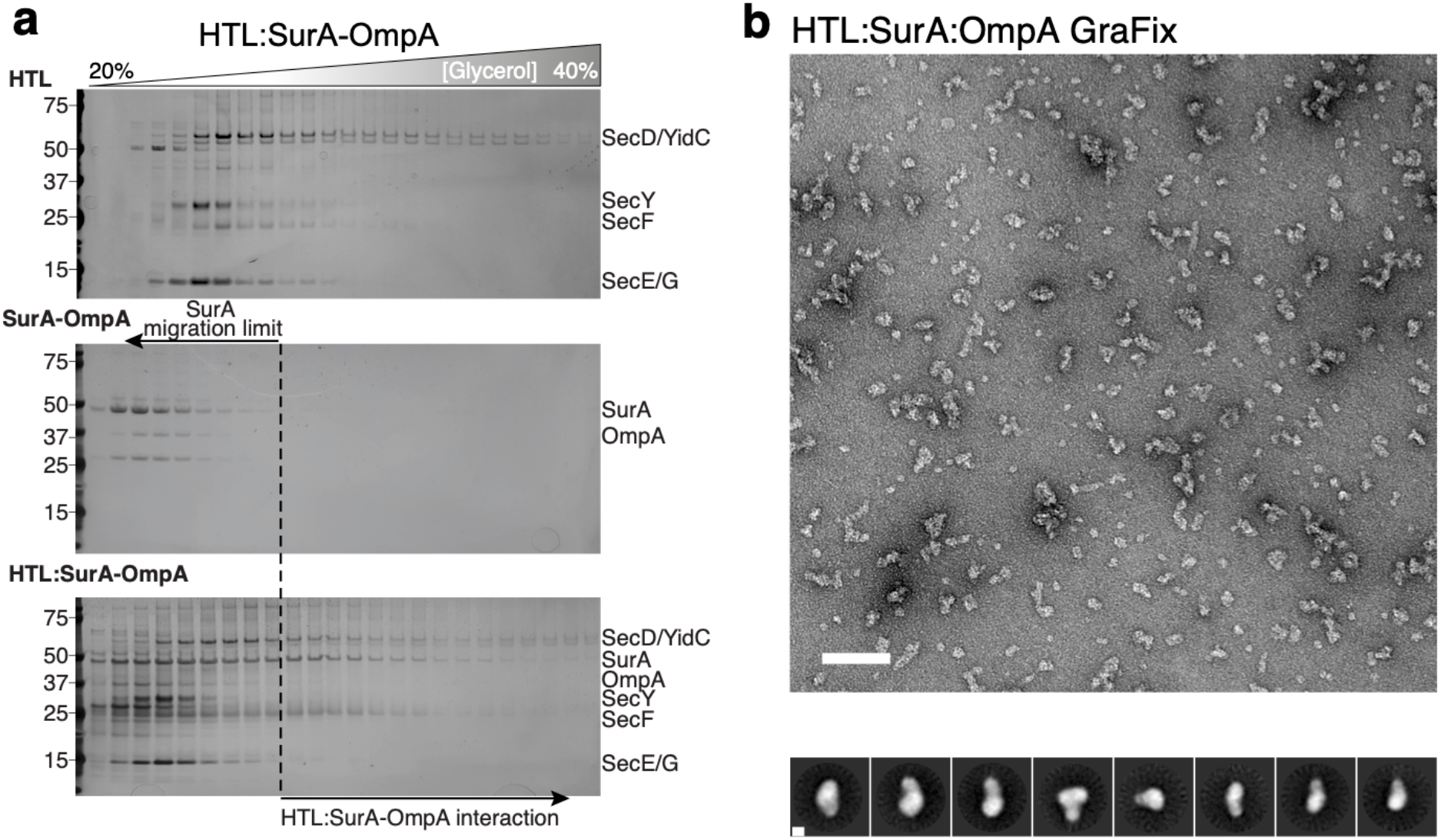
Negative stain EM micrograph and glycerol gradient centrifugation of HTL-SurA-OmpA. For **(a)**, the experiment was conducted as in Fig. S2. An EM field of HTL:SurA:OmpA prepared using GraFix is shown (**b**). Reference-free (RF) class averages of the biggest populations from the micrograph is shown (bottom). Micrograph scale bar, 1000 Å. RF scale bar, 100 Å.

